# Genetic and peripheral visual system changes underlie evolving butterfly mate preference

**DOI:** 10.1101/2022.04.25.489404

**Authors:** Nicholas W. VanKuren, Nathan P. Buerkle, Erica L. Westerman, Alexandria K. Im, Darli Massardo, Laura M. Southcott, Wei Lu, Stephanie E. Palmer, Marcus R. Kronforst

**Author notes:** Corresponding Authors SEP, MRK. These authors contributed equally to this work (Nicholas W. VanKuren, Nathan P. Buerkle). Department of Neuroscience, Yale University, New Haven, Connecticut 06510 (Nathan P. Buerkle). Department of Biological Sciences, University of Arkansas, Fayetteville, Arkansas 72701 (Erica L. Westerman).

## Abstract

Many studies have linked genetic variation to behavior, but less is known about how that variation alters the neural circuits that drive behavior. We investigated the genetic and neurobiological basis of courtship preference variation in *Heliconius* butterflies, which use vision to identify appropriate mates based on wing color patterns. We found that *Heliconius cydno* preference variation was strongly associated with genetic variation and differential expression of *senseless-2*, a gene predominantly expressed in the eye. Further measurements of photoreceptor sensitivities revealed differences in inter-photoreceptor inhibition of ultraviolet-sensitive cells corresponding to courtship preference variation. Our results reveal a genetic basis for preference/cue co-evolution, suggest a link between *sens-2* and visual system variation, and support the idea that changing peripheral neural computations can significantly alter essential behaviors.

**Summary:** Genetic and expression variation of *senseless-2* and inter-photoreceptor inhibition predict visual mate preference in a clade of diverse butterflies.

---

Behavioral evolution requires genetic variation that ultimately alters neural circuits. Many studies have mapped the genetic basis for behavioral evolution (*1*–*4*), but often missing is the connection to the neural changes that mediate these shifts in behavior. The peripheral nervous system appears to be an evolutionarily labile target for modification, presumably because changes in receptor sensitivity can avoid potentially deleterious effects associated with changing a complex central brain (*5*–*8*). However, simple shifts in receptor sensitivity may be insufficient to enact large behavioral changes, instead requiring more significant changes in how downstream circuits process sensory information (*9*–*12*). An integrated approach can reveal both where genetic effects are likely to take hold in the brain and how those changes affect neural computation.

The co-evolution of wing color and courtship preference in Neotropical *Heliconius* butterflies presents an excellent system to integrate genetic and neurobiological approaches to behavioral evolution. Visual perception of wings guides courtship behavior, as males preferentially court females with the same mimetic wing color pattern (*13*–*15*). Furthermore, the loci controlling color and preference in the *Heliconius cydno* clade are linked, a phenomenon known as genetic coupling (Fig. 1A; *16*). Males homozygous at the wing color locus tend to court females with the same color, while heterozygotes court both colors equally. We recently showed that *aristaless-1* controls the switch between dominant white and recessive yellow wing colors (*17, 18*), but the genetic variation underlying preference and its effects on neural circuits responsible for courtship behavior remained unknown. Here, we leverage preference variation in *Heliconius cydno* to identify a putative courtship gene, localize its expression to the eye, and find a difference in inter-photoreceptor connections that can potentially explain male preferences for white or yellow females.

**Figure 1.**
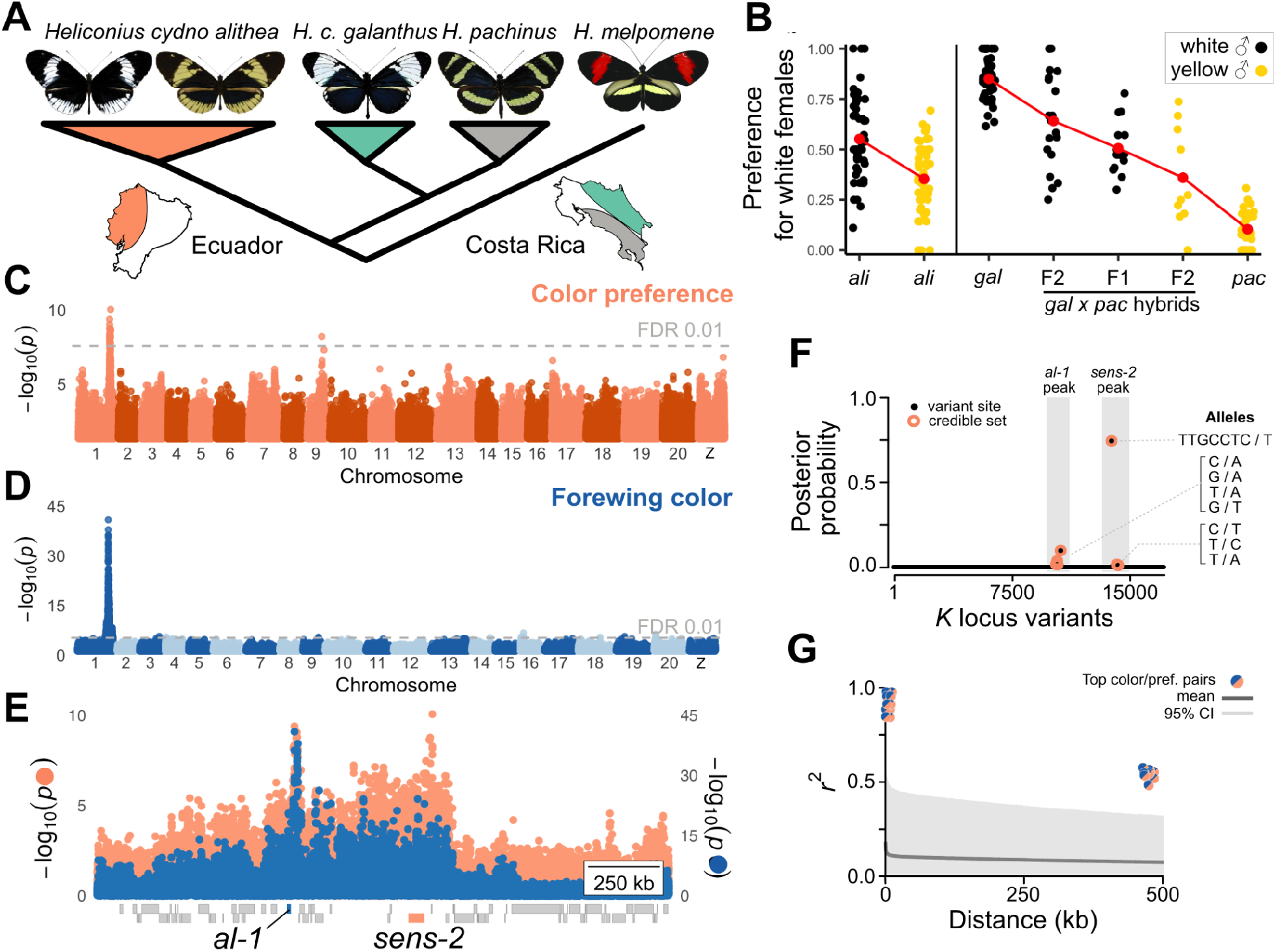
Genetic basis of *Heliconius cydno* preference variation. **A**, The *cydno* complex and its sister taxon, *H. melpomene*. **B**, Male mate preference, i.e. the proportion of courts directed at white females, from refs. (*14, 19*). Each point represents preference of one male; red points are means. **C-E**, GWA of *H. c. alithea* male mate preference and color genome-wide (**C**,**D**) and at the *K* locus (**E**). Boxes below the x-axis are gene models. **F**, Posterior inclusion probability for each variant in the *K* locus, calculated using *SuSiE-RSS (20)*. **G**, Pairwise linkage disequilibrium between top preference and color variants relative to empirical genome-wide levels.

We began characterizing the causes of preference differences using genome-wide association (GWA) in the polymorphic subspecies *Heliconius cydno alithea* (Fig 1; figs. S1, S2; Table S1;*19)*. GWA identified variants significantly associated with preference on two chromosomes, with the majority found in two narrow peaks in the *K* locus, a region which includes *al-1* and the previously identified preference quantitative trait locus (Fig. 1C; *14*). One peak co-localized with the known wing color locus, while the second, stronger peak was 482 kb away (Fig. 1C-E). Overall, *K* locus variants explained 20.5% of preference variance, consistent with the observed behavioral difference (Fig. 1B). Additionally, Bayesian analysis identified a credible set of eight putative causal sites in this region, with an indel in the second peak having the highest posterior probability (Fig. 1F; *20*). The top color-and preference-associated variants were in extremely high linkage disequilibrium (LD), but LD was not generally elevated across the *K* locus and we found no evidence for inversions or other genomic features to explain the high pairwise LD (figs. S4, S5). Rare recombination was indicated by mismatches between wing color and preference index (Fig. 1G, fig. S3).

The most significant preference variants were 25 kb upstream of the transcription factor gene *senseless-2* (Fig. 1E). While the function of *sens-2* is unknown, its paralog *senseless* is essential for peripheral sensory structure development in *Drosophila* (Fig. 2A; *21*–*23*). Consistent with a potential role in courtship behavior, we found differential *sens-2* expression between white *H. c. galanthus* and yellow *H. c. alithea* heads during mid-to late pupal stages, and subsequent tissue specific RNA-sequencing showed expression primarily in the eye (Fig. 2B; fig. S6). Furthermore, antibodies against Sens-2 specifically labeled nuclei of all photoreceptors, as well as the pigment cells that optically isolate neighboring ommatidia, in pupal and adult eyes (Fig. 2C-D, figs. S7, S8). We observed qualitatively similar expression patterns in all *H. c. galanthus* and *H. c. alithea*. In contrast to eyes, Sens-2 staining in the central brain was exclusively cytoplasmic and limited to the adult mushroom body (fig. S9).

**Figure 2.**
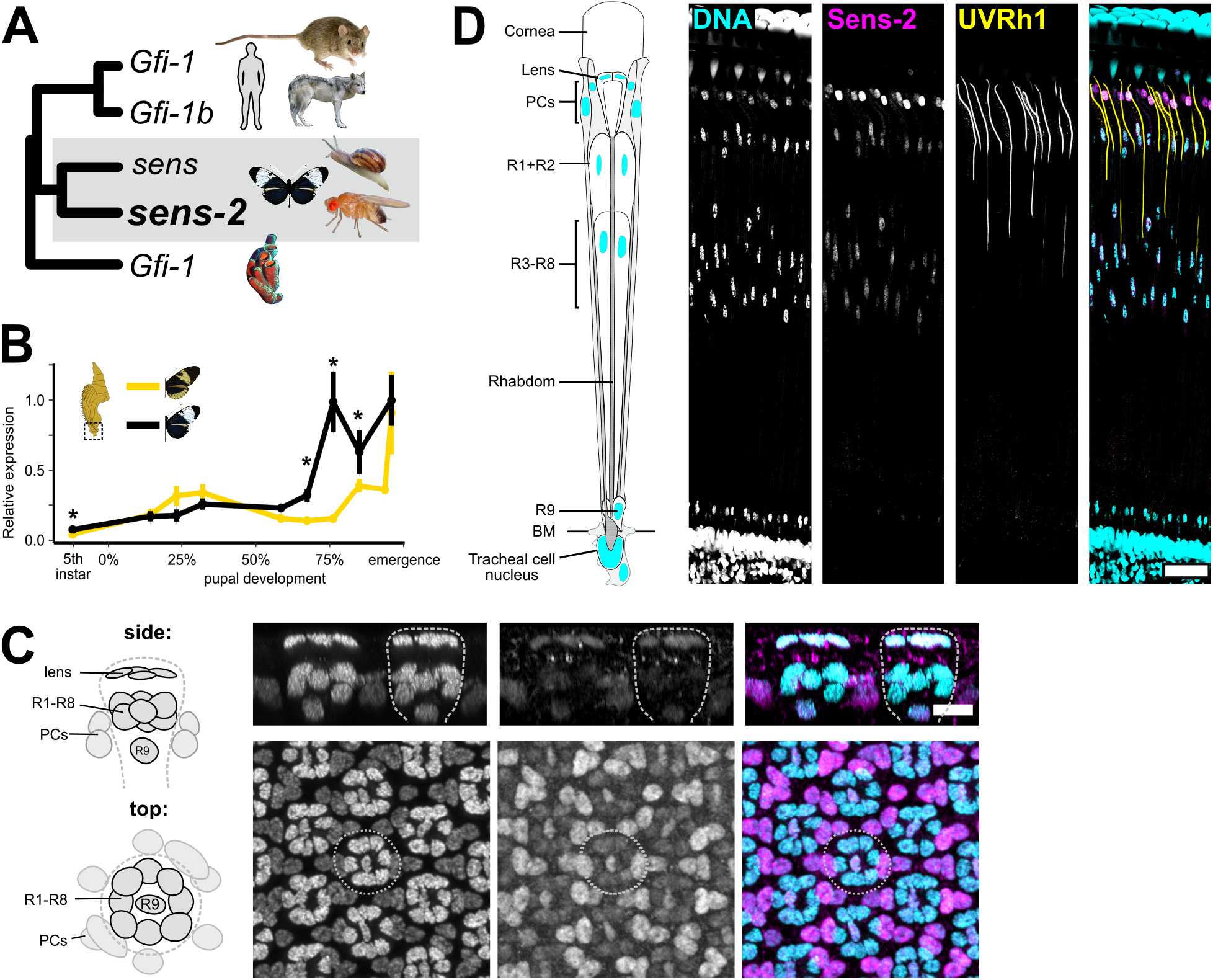
*senseless-2* evolution and expression in the *Heliconius* eye. **A**, Condensed neighbor-joining tree based on *sens-2* homologs identified by BLASTp (fig. S16). **B**, *sens-2* qPCR measurements in yellow *H. c. alithea* and white *H. c. galanthus* pupal heads. **C**, Anti-Sens-2 antibody staining in yellow *H. c. alithea* pupal retinas at ∼50% pupal development. Diagrams show nuclei; dashed lines outline single ommatidia; PC: pigment cell. **D**, Anti-Sens-2 antibody staining in adult white *H. c. alithea*. BM: basement membrane. Scale bars: **C** 10 μm, **D** 50 μm.

Our results suggest that variation in *sens-2* function may affect courtship behavior by altering the functional organization of the eye. We tested this hypothesis by recording from single photoreceptors for butterflies separated into six groups on the basis of species, wing color, and courtship preferences (Fig. 1A,B, Fig. 3). We first observed that every butterfly expressed a combination of UV-, blue-, green-, and red-sensitive photoreceptors (Fig. 3, fig. S10). Only UV cells showed a difference in spectral tuning across groups, with peak sensitivity varying continuously between 350 and 400 nm due to varying degrees of co-expression of an ancestral UV1 opsin and genus-specific UV2 opsin within single photoreceptors (Fig. 3A-C; *24, 25*).

**Figure 3:**
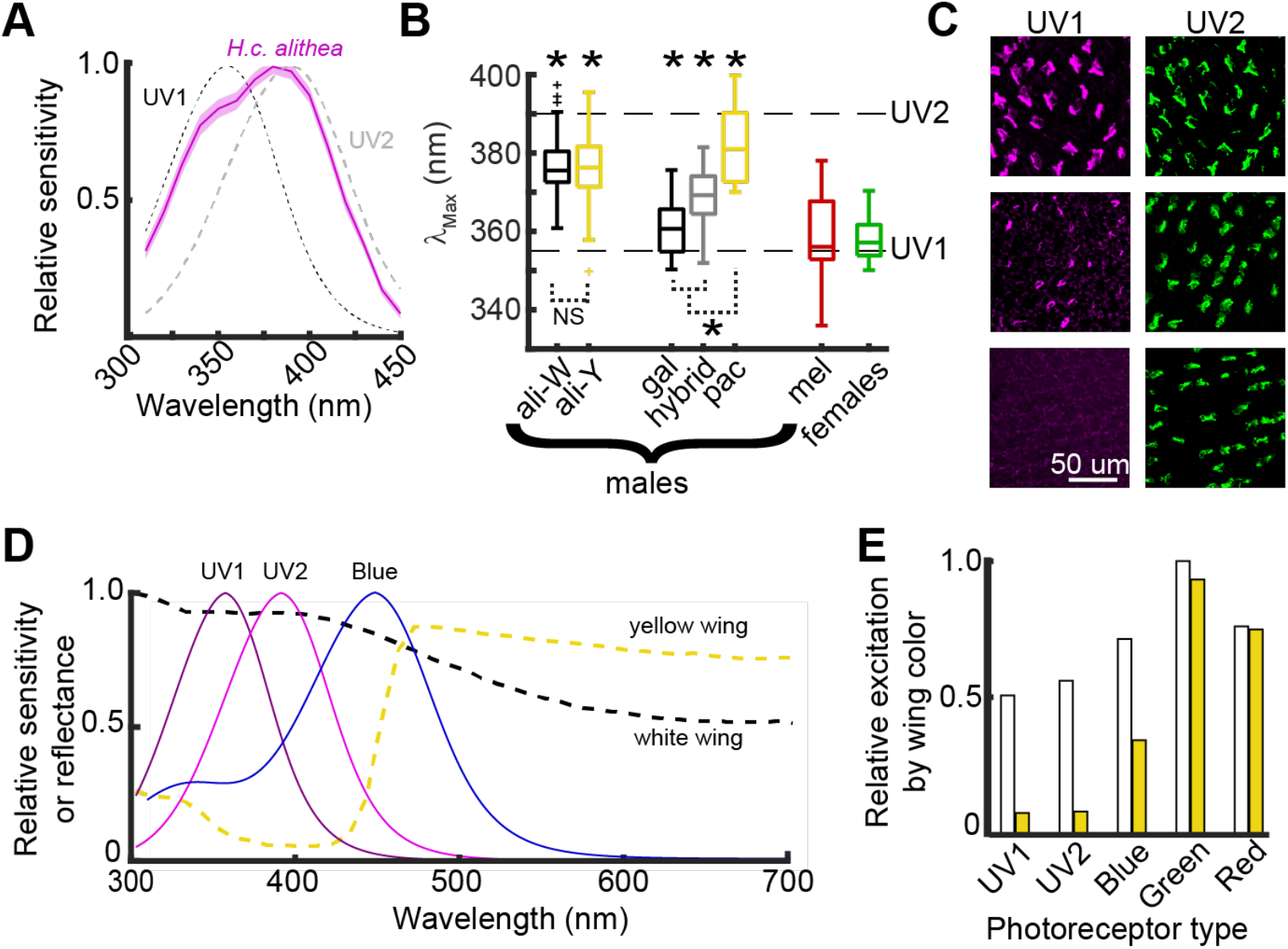
Variability in the spectral tuning of UV sensitive photoreceptors cannot explain courtship preference. **A**, Average tuning curve for all UV photoreceptors for *H. c. alithea* males. Shading shows the standard error. Dotted lines show the expected tuning of the ancestral UV1 opsin and derived, genus-specific UV2 opsin. **B**, Boxplots show the distribution of the wavelength of peak sensitivity for UV photoreceptors. Asterisks above indicate groups that are significantly different from the expected tuning of both UV1 and UV2 (one sample t-tests, p < 0.05 after Holm-Bonferroni correction). Spectral tuning varied between groups (F_6,168_ = 21.70, p < 0.001), with the asterisk below indicating significance (t-test, p < 0.01 with Holm-Bonferroni correction; n = 43, 40, 18, 19, 8, 25, 22). **C**, Cross-sections of the eye immunostained for UV1 and UV2 opsins. Depicted are examples for three *H. c. alithea* males with strong UV2 expression and variable degrees of UV1 expression. **D**, Spectral sensitivity curves overlayed with the spectral reflectance of white and yellow wings. **E**, Normalized convolution between photoreceptor spectral tuning and wing reflectance shows the relative excitation of each cell type for white and yellow wings.

Comparing across groups, UV photoreceptor tuning exhibited a weak correlation between UV2 expression and preference for yellow females, as well as a clear sexual dimorphism where all females expressed only UV1 (Fig. 3B). However, this relationship cannot explain courtship preference variation for two main reasons. First, UV tuning was not significantly different between white and yellow *H. c. alithea* despite their behavioral differences (Fig. 3B). Second, white and yellow wings primarily differ by the presence and absence of UV reflectance, respectively (Fig. 3D). Thus, white wings strongly excite and yellow wings weakly excite UV cells regardless of the specific wavelength of peak sensitivity (Fig. 3E). Instead, stronger UV2 in males that will court yellow females is consistent with data showing that UV2 enhances the discriminability of a genus specific yellow pigment from the yellow pigments of distantly related species that mimic *Heliconius* wing patterns (*26*).

More generally, this result indicates that simple changes in receptor sensitivity cannot explain preference, instead requiring a more significant change in how visual information is processed (Buerkle et al., unpublished). Closer inspection of our recordings revealed a physiological signature of inhibition between photoreceptors (as observed in other species; *27*– *30*) that could potentially provide this larger computational change (Fig. 4). Insect photoreceptors depolarize in response to light and use histamine as their inhibitory neurotransmitter (*29, 31*). However, we measured unexpected hyperpolarizing responses to long wavelength stimuli in a subset of UV and blue sensitive photoreceptors (Fig. 4A,C, fig. S11,12).

**Figure 4:**
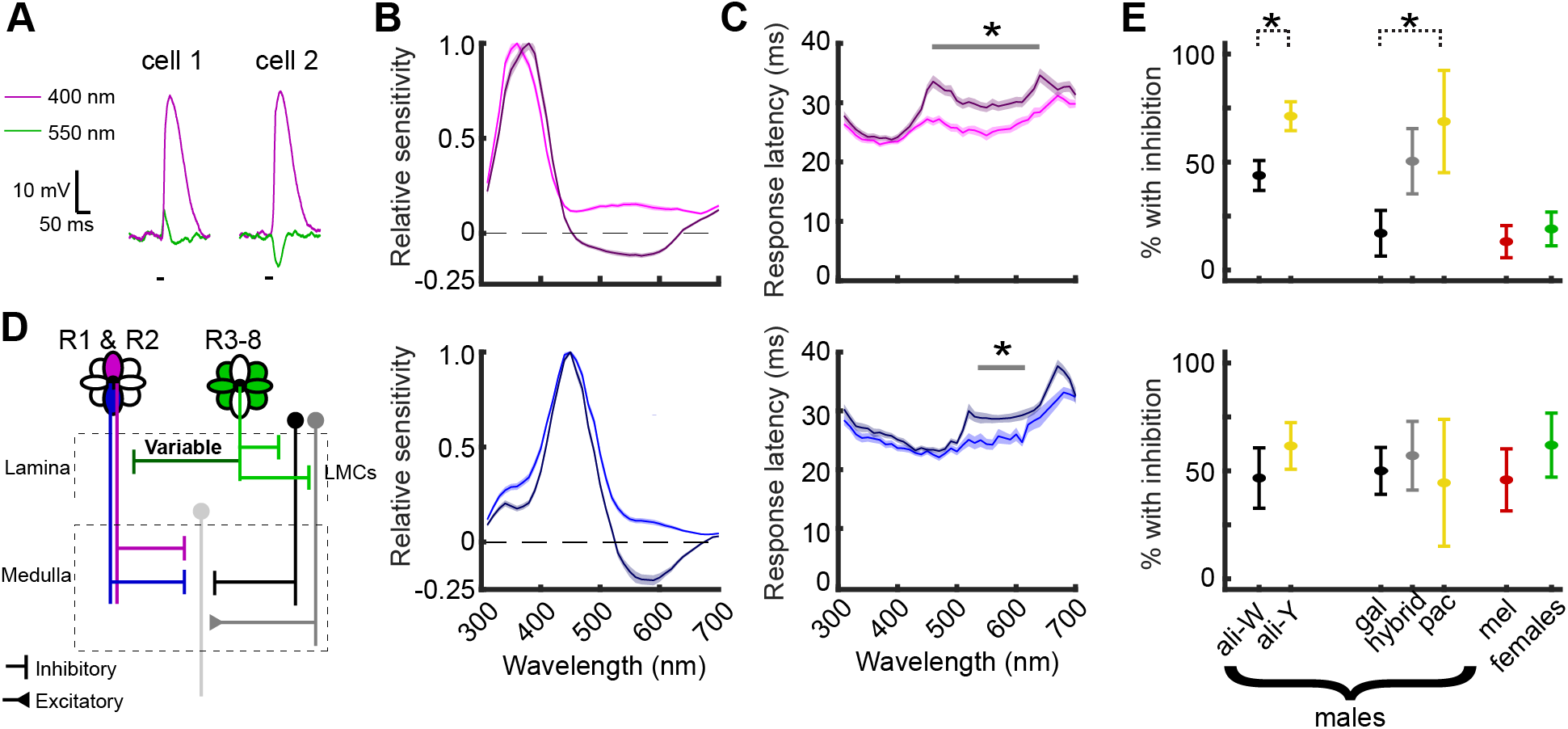
Lateral inhibition onto UV photoreceptors correlates with courtship preference. **A**, Traces show the responses of two UV photoreceptors for *H. c. alithea* males. **B**, Tuning curves for UV (top) and blue (bottom) photoreceptors separated into groups that either had (n = 76 UV, 59 blue) or did not have (n = 104, 67) hyperpolarizing responses. Shading shows the standard error. **C**, Response latency for the cells in panel B. Asterisks indicate significant differences (paired t-tests, p<0.05 with Holm-Bonferroni correction). **D**, Schematic showing lateral inhibition from R3-8 photoreceptors as the source of hyperpolarization. LMC: lamina monopolar cell **E**, Proportion of UV (top) and blue (bottom) cells with inhibitory inputs. Estimated for each butterfly (n = 13, 14, 10, 9, 4, 16, 10; 1-6 cells per butterfly), and showing the mean ± SEM. Groups were compared with an ANOVA (F_6,69_=7.35, p<0.001) and t-tests (p=0.001 for *H. c. alithea* comparison and p=0.037 for *H. c. galanthus* vs. *H. pachinus*). Both comparisons were also significant when comparing total count (*H. c*. alithea: *X*^2^=4.7, p=0.03; *H. c. galanthus* vs. *H. pachinus*: *X*^2^=8.3, p=0.004). Total cells for UV = 43, 40, 18, 19, 8, 30, 22; Blue = 21, 30, 22, 21, 5, 15, 12.

Photoreceptor response properties indicated that hyperpolarizing responses originated from green or red cells making inhibitory synapses onto UV and blue cells (Fig. 4B). First, hyperpolarizing responses were delayed by 5.5 +/-3.6 ms relative to depolarizing responses, consistent with monosynaptic inhibition (Fig. 4D, fig. S13). Second, we recorded from UV cells in the presence of an 534 nm LED that strongly excites green photoreceptors (fig. S14). Consistent with the green light inducing a persistent lateral inhibitory current, the resting potential of UV cells with hyperpolarization decreased by 5.5 +/- 4.8 mV, while cells without hyperpolarization were unaffected. Additional differential effects of the LED on UV-cell response magnitude and temporal profile points to the existence of distinct photoreceptor subtypes. Together, these results indicated that UV and blue photoreceptors have variable anatomical connections, with some that could support important downstream computations like color-opponency.

Comparing inter-photoreceptor connections across groups showed that differences in UV inhibition, but not blue, aligned with differences in courtship preferences (Fig. 4E). We again observed a sexual dimorphism, with UV photoreceptors for all females and *H. melpomene* rarely showing evidence of inhibition. For males that prefer yellow females (*H. pachinus* and yellow *H. c. alithea*), 70.8% of UV photoreceptors had inhibitory responses, while only 16.7% had inhibition for white-preferring males (*H. c. galanthus*). Males with no preference (white *H. c. alithea* and *H. c. galanthus* X *H. pachinus* hybrid offspring) were intermediate, with 46.8% of UV cells having inhibitory responses. This similarity is consistent with post-hoc genetic analysis showing that 91% of white *H. c. alithea* males were heterozygous at the *K* locus. Further, the significant difference between white and yellow *H. c. alithea* was particularly striking because the *K l*ocus is the only differentiated region of the genome between these two groups with different preferences (fig. S15). Finally, s*ens-2* expression data matched the timeframe when photoreceptors form synapses (*32*), with higher expression in *H. c. galanthus* supporting a role for it in repressing the formation of synapses onto UV cells.

Our working hypothesis for how these connectivity differences drive courtship preference proposes that UV stimuli promotes approach behavior, and inhibition functions as a gain control that titrates the propagation of these signals into courtship circuitry. Yellow is the ancestral *H. cydno* wing color (*17*), so rather than a *de novo* choice between white and yellow females, the presence of UV inhibition is presumably part of making yellow the default color preference. Assuming UV light has a positive courtship valence, removal of inhibition when white wings evolved would allow the strong UV component of white wing reflectance to override the default preference, with shared opponent computations or homeostatic compensation suppressing the white male’s approach towards yellow females.

The evolvability of the periphery typically refers to changes in receptor sensitivity like what we observed in the spectral tuning of UV photoreceptors (Fig. 3), while larger computational changes are thought to require changes to complex central circuits. Our work merges these ideas, suggesting a role for inter-photoreceptor inhibition in white vs. yellow preference (Fig. 4), and revealing that the circuit architecture of the periphery may also be subject to rapid evolution. These lateral connections, reminiscent of horizontal cells in the vertebrate retina, have now been observed in *Drosophila* and more than ten butterflies. Thus, our results may be broadly applicable to other systems and sensory modalities.

Overall, our integrated genetic and neurobiological approaches provided us with a fuller picture of the mechanisms that drive/underlie co-evolution of wing color and courtship preference. Co-evolution between color and preference in *H. cydno* is mediated by genetic coupling between two physically separate but linked loci rather than a single pleiotropic gene or genome structural variation. Theory predicts that speciation should be rare when preference and cue are controlled by separate loci because recombination should quickly break down association between the two traits (*33*–*35*). However, this mechanism of coupling is common; this and several recent studies have found preference/cue coevolution mediated by separate but linked QTL (*36, 37*). Coupling may be partly caused by assortative mating itself (*38*) but is likely enhanced in *Heliconius* by natural selection against locally rare aposematic wing colors, which eliminates individuals with mismatched color and preference alleles (*39*). *Heliconius* have rapidly evolved myriad wing color patterns, and genetic coupling should entail similarly rapid adaptations of the nervous system. Thus, genes important to the functional organization of an evolutionarily labile periphery may play an important role in facilitating the initial stages of the speciation process.

## Acknowledgements

We thank Nicola Chamberlain, Ryan Hill, Durrell Kapan, Lawrence Gilbert, Jacob Olander, Andres Vega, Sumitha Nallu, and Anastasia Weger for helping collect data and for valuable discussions, and Michael Perry for providing antibodies and discussions. We thank Michiyo Kinoshita and Kentaro Arikawa for hosting NPB and providing training on physiology methods.

## Author contributions

NWV and NPB designed experiments, performed experiments, analyzed data, and wrote the paper. ELW performed experiments and analyzed data. AKI and DM performed experiments. WL analyzed data. SEP and MRK designed experiments, analyzed data, acquired funding, and wrote the paper.

## Data and materials availability

Sequencing data are publicly available through NCBI, BioProjects PRJNA802828 and PRJNA802836. Sens-2 antibody available upon request.

## Supplementary materials

Materials and methods

Table S1-S7

Fig S1-S17

References (40-74)

