## Supplementary Materials for "Genetic and peripheral visual system changes underlie evolving butterfly mate preference"

### Materials and Methods

#### Animals

The animals used here comprise butterflies from four different taxa. For the GWA, *H.c. alithea* males that were previously tested for courtship behavior were used (*19*). For all other experiments, butterflies were housed in a greenhouse breeding colony at the University of Chicago that was regularly supplemented with new individuals. *H.c. galanthus* and *H. melpomene* pupae were obtained from breeders in Costa Rica, and *H.c. alithea* from breeders in Ecuador. *H. pachinus* and F1 *H.c. galanthus* X *H. pachinus* hybrids were bred in Panama and adults were transported to Chicago for experiments. Collection, rearing, import and export were done under permits from Ecuador, Panama, Costa Rica, and USA.

#### *Heliconius cydno galanthus* genome assembly and annotation

We isolated DNA from thorax of a single adult *H. cydno galanthus* female using phenol-chloroform extraction. We constructed Illumina paired-end (PE) libraries with insert sizes 250 bp and 500 bp using the KAPA Hyper Prep Kit (KR0961) and 1 µg genomic DNA each. We constructed mate pair (MP) libraries with insert sizes of 2 kb, 6 kb, and 15 kb using the Nextera Mate Pair Library Prep kit (FC-132-1001) and 4 µg genomic DNA each. We pooled these libraries in a ratio of 50:18:10:17:4 and sequenced them 2 x 100 bp on a single lane of Illumina HiSeq 4000. We performed additional 2 x 100 bp HiSeq 4000 sequencing on the PE libraries. Raw sequencing data is available in the NCBI SRA (Table S3). We trimmed low-quality regions and remaining adapters from raw PE reads using Trimmomatic v0.36 (*40*) and from MP libraries using the platanus_internal_trim tool from Platanus v1.2.4 (*41*). Trimmed libraries were assembled using Platanus v1.2.4 (default settings) and the assembly polished using Redundans v0.13a (default settings) (*42*). For plotting, we mapped *H. cydno* scaffolds to the *H. melpomene* v2.5 reference assembly using Minimap2 and RagTag, which allowed us to identify and split chimeric *H. cydno* scaffolds and assign *H. cydno* scaffolds to *H. melpomene* chromosomes (*43*, *44*). Mapping was only taken into account when plotting results.

We annotated *H. cydno* scaffolds using EvidenceModeler 1.1.1 (*45*). We first assembled the *H. cydno* transcriptome *de novo* using RNA-seq data generated by Walters et al. (*46*), Nallu et al. (*47*), Rossi et al. (*48*), and newly-generated data for eyes and brains (see below) using Trinity v2.10.0 (*49*). RNA-seq data was also mapped to *H. cydno* scaffolds using STAR 2.6.1d (*50*), and the resulting alignments used to generate genome-guided assemblies using Trinity and StringTie 1.3.1 (*51*). We combined *de novo* and genome-guided assemblies using PASA (*52*). Evidence for protein-coding regions came from mapping the UniProt/Swiss-Prot (2020_06) database and all Papilionoidea proteins available in NCBI's GenBank nr protein database (downloaded 6/2020) using exonerate (*53*). We identified high-quality multi-exon protein-coding PASA transcripts using TransDecoder (transdecoder.github.io), then used these models to train and run Genemark-ET 4 (*54*) and GlimmerHMM 3.0.4 (*55*). We also predicted gene models using Augustus 3.3.2 (*56*), the supplied *heliconius_melpomene1* parameter set, and hints derived from RNA-seq and protein mapping above. Augustus predictions with >90% of their length covered by hints were considered high-quality models. Transcript, protein, and *ab initio* data were integrated using EVM with the weights in Table S4.

Raw EVM models were then updated twice using PASA to add UTRs and identify alternative transcripts. Gene models derived from transposable element proteins were identified using BLASTp and removed from the annotation set. The final annotation comprises 16,788 genes encoding 33,051 protein-coding transcripts.

*Heliconius cydno alithea* genome re-sequencing and variant calling

Genomic DNA was isolated from thorax of 113 *H. c. alithea* males studied by Chamberlain et al. (*19*) using chloroform extractions (Table S1). Illumina paired-end libraries were constructed using the KAPA Hyper Prep Kit (KAPA Biosystems) or Nextera Library Prep Kit and sequenced to ~15X using PE100 bp on an Illumina HiSeq2500 or 4000 at the University of Chicago Functional Genomics Facility.

Low-quality regions and adapters were trimmed from raw reads using Trimmomatic before mapping to the *H. c. galanthus* reference using bowtie2 v2.3.2 with default settings except --very-sensitive-local (*57*). We then marked PCR duplicate reads with Picard and realigned around putative indels using the Genome Analysis Toolkit (GATK) 3.8 (*58*, *59*). SNP and indel calling was performed using the GATK's UnifiedGenotyper module with the heterozygosity priors set to 0.02 and 0.0002 for SNPs and indels, respectively. We merged SNPs and indels, then filtered out sites with >10% missingness and minor allele frequency <0.05 before read-backed phasing and imputation with SHAPEIT2 (*60*). The final variant call set comprised 7.39 million SNPs and 1.56 million indels.

Genomics analyses

We identified global covariates using principal components analysis (PCA; Fig S1). We first pruned our final variant set using PLINK's (1.90) linkage disequilibrium-based pruning ( i.e. --*indep-pairwise 1000 100 0.80*) (*61*), leaving 4.8 million variants in PCA.

To estimate allele frequency divergence between yellow and white *H. c. alithea,* we calculated genome-wide $F_{ST}$ in 10 kb (not shown) and 50 kb (Fig S13) non-overlapping windows using VCFtools 0.19 (*62*) and Weir and Cockerham's method (*63*).

LD and searches for recombination suppression mechanisms

We calculated genome-wide linkage disequilibrium (LD) decay using a random sample of 600 million pairs of variants and PLINK 1.90, then summarized and plotted the results in R. Pairwise LD between the top 10 color and preference variants was also calculated using PLINK 1.90.

We searched for evidence of genome structural variation in the *K* locus using read mapping methods implemented in delly v0.7.6 (*64*) and pindel (*65*). These methods search for large (>50 bp) insertions, deletions, inversions, and translocations based on read pair mapping orientations. To maximize sensitivity, we called SVs in each sample separately using lenient filters (read mapping quality > 0, minimum one read pair supporting an SV), merged calls of the same type using delly, then jointly genotyped all samples for all SVs using delly.

Recombination rates are negatively correlated with the density of repeat elements in a variety of organisms (*66*). We tested if increased pairwise LD between the color and preference GWA peaks may be influenced by repeat density in the *K* locus by calculating the density of repeat elements across the locus using RepeatMasker predictions. We found no evidence for increased repeat density between the color and preference peaks, and somewhat decreased density around *K* relative to genome-wide levels (Fig S5). These figures display the proportion of masked sequence in 50 kb (genome-wide) or 10 kb (K locus) windows. This result did not change when we limit to just putative TEs.

Genome-wide association analyses

Details on the collection and initial analyses of the male courtship data are found in Chamberlain et al. (2009). We performed additional analyses based on the subset of males that we re-sequenced. We found a strong correlation between a male's lifetime preference (i.e. the proportion of courtship events directed at white-winged females) and the total number of courts that male performed (Fig S2), such that males of both colors eventually make, on average, random mate choices. We hypothesize that this is due to males becoming frustrated after a certain number of thwarted mating attempts - males were captured after courting, then released back into the testing cage, so they were never allowed to copulate (*19*). Male preference began to regress after males reached about 20 courts (Fig S2C), so we limited our analyses to preference estimates based on the first 20 courts for each male.

We included males with a single court because a male's first court is significantly predictive of lifetime preference, even when controlling for male wing color.

*lm(preference ~ first_court + male.fwc, data = x)*

| | **Estimate** | **S.E.** | ***t*** | **Pr(>\|*t*\|)** |
|---|---|---|---|---|
| **(intercept)** | 0.742 | 0.040 | 18.563 | <2e-16 |
| **First court** | -0.316 | 0.053 | -5.979 | 6.39e-8 |
| **Male color** | -0.223 | 0.053 | -4.231 | 6.29e-5 |

We performed GWA separately for preference and forewing color using GEMMA 0.98 (*67*). The final mixed model included genetic relatedness (calculated using GEMMA) and the first three PCs from the analysis described above, which each explained >6% total variance. The final preference model was therefore:

$$preference \sim genotype + PC1 + PC2 + PC3 + GRM + \varepsilon$$

We plotted the results, using likelihood ratio test *p*-values, using the gwplotting R package (https://github.com/nwvankuren/gwplotting.git).

SuSiE-RSS analysis

For statistical fine-mapping using SuSiE-RSS (*20*, *68*), we selected the 17,228 SNPs, spanning from the *K* locus and surrounding regions chromosome 1 (pictured in Fig. 1E). The input z-scores are calculated from the estimates of marginal association for each variant site divided by corresponding standard errors (available in GEMMA's output). The LD matrix is calculated by PLINK. The maximum allowed causal variant number was set at 10 and the default sparse prior is used.

qPCR

Staged pupal *H. c. galanthus* and yellow *H. c. alithea* heads were collected and stored in TRIzol (Ambion) at -20°C until RNA extraction. We synthesized cDNA using the ABI High Capacity cDNA Synthesis Kit and 1 ug total RNA for each sample. Samples were diluted 1:10 in TE before using 1 uL in qPCRs assaying *sens-2* and *elongation factor 1-alpha* expression in each sample (Table S5). *Sens-2* expression levels were calculated relative to *EF1a* as $2^{(CT,EFa\ -\ CT,sens-2)}$ and scaled to the highest value before plotting.

RNA-seq

We performed RNA-seq on retinas and central brains of *H. c. galanthus* males and females at 2, 4, and 7 days after pupa formation. Tissues were dissected in ice cold PBS, then immediately stored in RNAlater (Ambion, USA) at -80°C until RNA extraction using TRIzol (Ambion, USA). The University of Chicago Functional Genomics Facility constructed libraries using Illumina TruSeq Stranded mRNA kits and sequenced libraries PE100 on an Illumina NovaSeq 6000. We quantified gene expression using the *H. c. galanthus* reference transcripts, salmon v1.4.0 (*69*) and tximport (*70*).

Antibody production and staining

We contracted GenScript (NJ, USA) to raise a new polyclonal antibody against the Sens-2 peptide antigen PELEVDSPPSSPRR in rabbit. Pupal retinas were dissected out in ice cold PBS, then fixed for 15 min in 4% formaldehyde in PBS at RT. Fixed tissues were then rinsed 3 x quickly and 4 x 15 min in PBST (0.3% Triton X-100 in PBS) before blocking for 1 hour in 5% normal goat serum (NGS) in PBST. Tissues were stained in 2% NGS/PBST with appropriate antibodies overnight at RT, washed 4 x 15 min in PBST, then stained with secondary antibodies in 2% NGS/PBST for 2 hr at RT before final washes of 3 x quick, 4 x 15 min, and 2 x 1 hr in PBST. Retinas were resuspended in Vectashield (Vector Laboratories, USA) overnight, then mounted in fresh Vectashield.

Adult eye and brain histology was performed as previously described (*25*). We removed cuticle, tracheae, and other non-neural tissue from around the brain and eyes in ice cold PBS, then fixed brain/eyes overnight in pre-chilled 4% formaldehyde at 4°C. Tissues were washed 3 x 15 min in PBS, 1 hr in 25% sucrose in PBS, and then fresh 25% sucrose in PBS at 4°C until sectioning (1 - 4 days). Tissues were mounted in OCT, sectioned anterior-posterior at 50 - 100 µm on a cryotome, arranged on Fisherbrand SuperFrost PLUS slides (Thermo Fisher, USA), and dried for 10 min on a 37°C slide warmer before proceeding immediately to washing and staining. We only used adults that had emerged within the previous 16 hours. However, adults >1 hour post-emergence had variable but high levels of off-target signal in tracheoles for DNA stains and most antibodies.

Antibodies were used at the following concentrations: rabbit α-sens-2 (1:1000 for IF, 1:200 for IHC), guinea pig α-UV1 (1:300 for IHC; Buerkle et al. *bioRxiv*), rat α-Pros (kindly provided by Mike Perry, UCSD; 1:200 for IF, 1:50 for IHC), mouse α-synapsin (Developmental Studies Hybridoma Bank 3C11; 1:40 for IHC), goat α-rat AlexaFluor 555 (Abcam 150158; 1:1000), goat α-mouse AlexaFluor 555 (Abcam 150114; 1:1000), goat α-rabbit AlexaFluor 488 (Abcam 150077; 1:1000), Hoechst 33342 (Invitrogen, USA H1399; 1:1000).

Samples were imaged on a Zeiss LSM 710 confocal microscope in the University of Chicago Department of Organismal Biology and Anatomy's Confocal Digital Imaging Facility. Images were processed in FIJI.

Sens-2 CRISPR/Cas9 knockouts

We attempted to knock out *sens-2* by using CRISPR/Cas9 to induce small frameshifts near the start codon to test the specificity of our α-Sens-2 antibody (Tables S5, S6). We identified CRISPR target sites within *sens-2* exon 1 using IDT's (USA) crRNA Design Tool, then synthesized sgRNAs and performed injections following Perry et al. (*71*). We screened for mosaic mutants and tested the specificity of our α-Sens-2 antibody by dissecting and staining ~50% pupal development retinas as detailed in the previous section. Initial high-resolution melt analyses on eggs two days after injection suggested high mutation efficiencies (30% - 100%), but hatching rates were consistently low (<10%) and survival to pupation even lower (<5%). These rates were consistent across a number of gRNAs, combinations of gRNAs, and concentrations of RNP (Table S6). Of the 17 pupae that we dissected and stained retinas for Sens-2, we observed four (23%) with clearly mosaic anti-Sens-2 staining patterns (Fig S15). Constitutive *sens-2* RNAi in *Drosophila melanogaster* is completely lethal, suggesting that *sens-2* is essential; we suspect that *sens-2* is also essential in butterfly development. Ideally we would test the effects of *sens-2* knockouts on courtship behavior, but the extremely low efficiency, sparse mosaicism, and difficulty generating homozygous *sens-2* mutant butterflies currently prevents such a direct test. Future work will explore alternative approaches to studying altered or loss-of *sens-2* function in these non-model organisms.

Intracellular Electrophysiology

For *in vivo* recordings, butterflies were restrained in a custom built collar with heated beeswax. A small hole was cut in the dorsal eye to allow for electrode penetration along the dorsal-ventral axis of the eye and covered with silicone grease to prevent dessication. A second small hole was cut near the mouthparts and a silver-chloride reference electrode was placed into the anterior portion of the head. The butterfly was then placed on a stage with the eye at the center of a Cardan arm perimeter device to allow for equivalent light stimulation at any spatial location.

Photoreceptor responses were evoked using monochromatic stimuli ranging from 310 nm to 700 nm in 10 nm increments. The light source was a dual Halogen-Deuterium lamp (DH-2000s, Ocean Optics), which was connected to a scanning monochromator (Monoscan-2000, Ocean Optics). Stimulus timing was controlled with an optical shutter (OZ Optics) and focused onto the butterfly eye using a collimator and lens (Edmund Optics). Every component was connected to each other using 1 mm fiber optic cables. Stimulus intensity was calibrated with a photodiode (Newport) and set to $1.5 \times 10^{15}$ photons/cm$^2$/s using a variable neutral density filter in a rotational motor (Newport). Three wavelengths (580, 650, and 660 nm) were excluded from the stimulation protocol due to artifacts from the deuterium bulb causing high and unstable intensities. Recordings were amplified with a 0.1X headstage and high impedance amplifier (AxoClamp 900A, Molecular Devices) and digitized at 10 kHz (DigiData1550, Molecular Devices).

Photoreceptors were recorded intracellularly using sharp electrodes made from borosilicate glass on an electrode puller (P-97, Sutter Instruments). Electrodes were pulled to a resistance between 90 and 120 MΩ and filled with 3 M KCl. Recordings were made exclusively from cells in the ventral half of the eye. All photoreceptors responded to white light with depolarizations of at least 30 mV. Stimuli were presented in a random order with 4 repeats per stimulus. Typically, responses were recorded at multiple intensity levels using neutral density filters (Thorlabs). After recording spectral responses, we also presented the wavelength that evoked the maximum response at 9 intensity levels that varied over 4 log units of attenuation. These V-Log(I) curves were used to transform the isoquantal spectral responses of the photoreceptors to a spectral sensitivity curve using the Naka-Rushton equation (*72*). The wavelength of peak sensitivity was estimated for each cell by fitting its responses with a standard rhodopsin tuning template (*73*). To measure response latency, we first measured the mean and standard deviation of the resting potential for 500 ms before the light flash. Onset latency was defined as the time for the response to exceed five times the standard deviation of this baseline.

For experiments with the LED, we used green LEDs with peak tuning at 534 nm and a full width half maximum of 12 nm. Six LEDs were attached to the monochromatic source and had an intensity of 3.2 X $10^{15}$ photons/$cm^2$/s. Spectral responses were recorded from each cell before, during, and after turning on the LEDs. This intensity did not bleach photoreceptor responses, as the full response magnitude was typically recovered within seconds of turning off the LED. Photoreceptors that did not recover at least 80% of the original response were discarded.

### Supplementary Figures

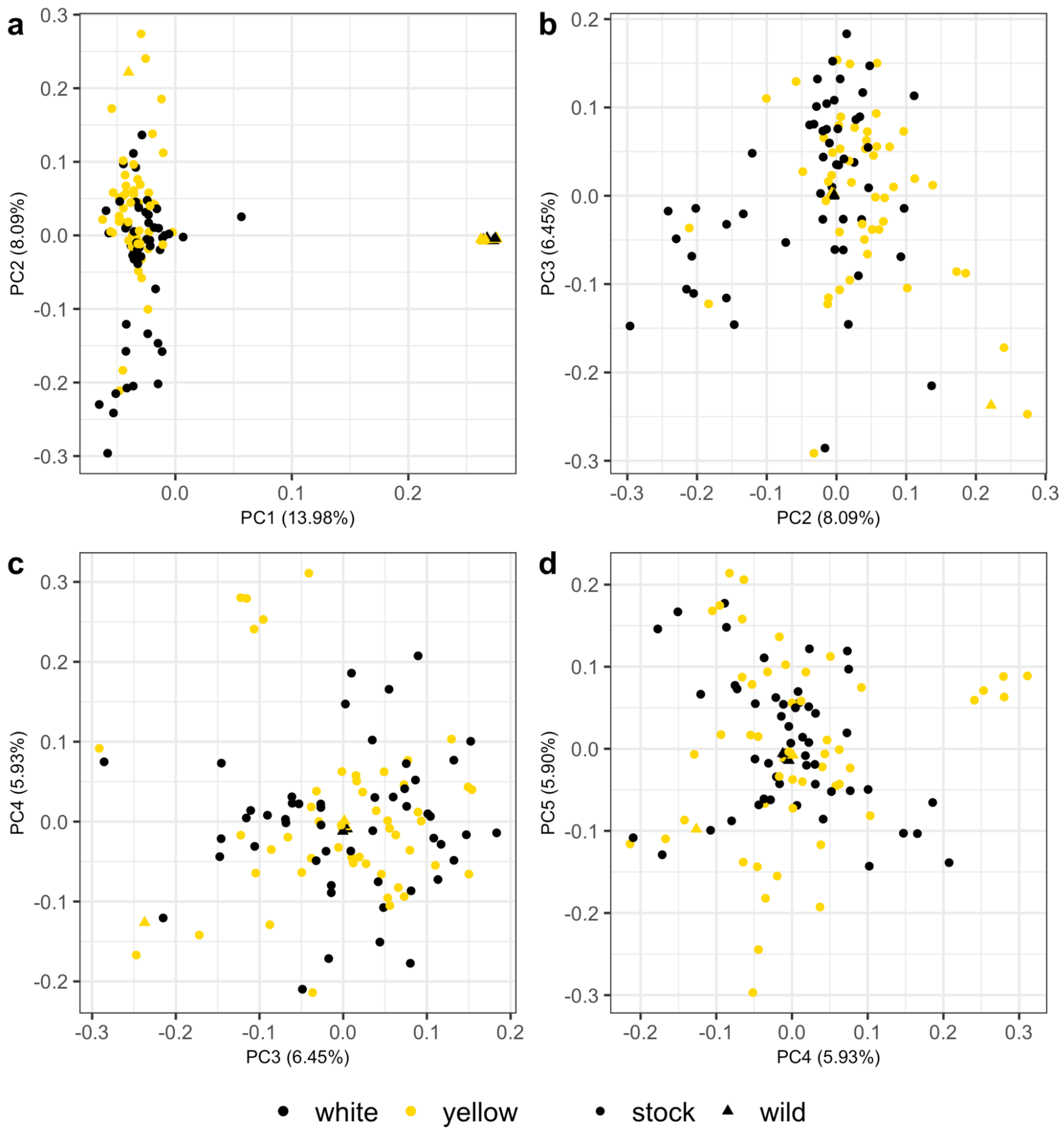


**Figure S1. Principal components analysis (PCA) of *H. c. alithea* genotype data.** PCs explaining >6% of variance were included as covariates in genome-wide association models.

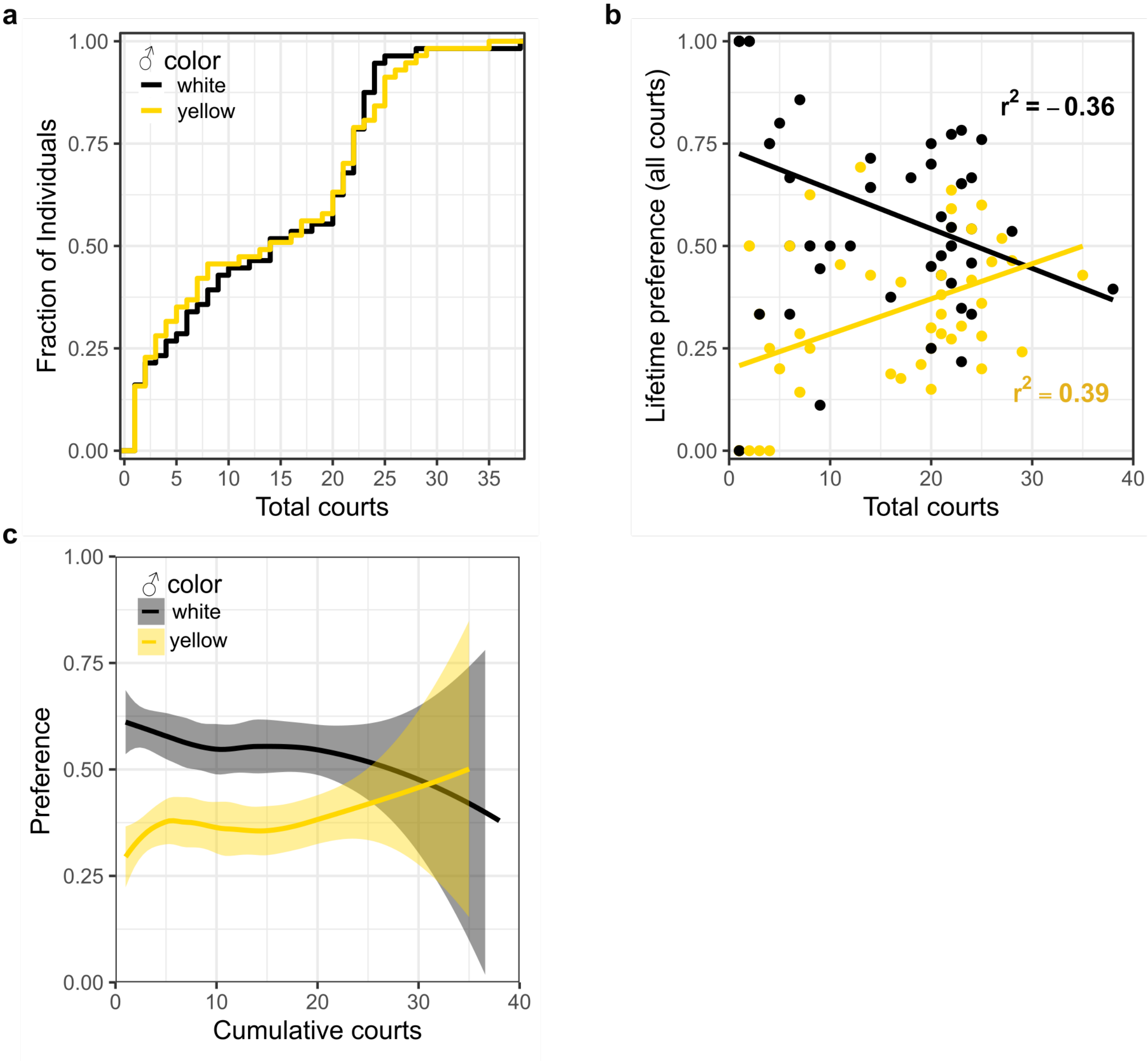


**Figure S2. *Heliconius cydno alithea* male preference data. a,** Cumulative distribution of courtship events in the Chamberlain data. **b,** Relationship between a male's total courts and his lifetime preference (i.e. the proportion of courts directed at white females). Each dot represents one male. **c,** Preference as a function of total courts. Shaded regions are standard errors.

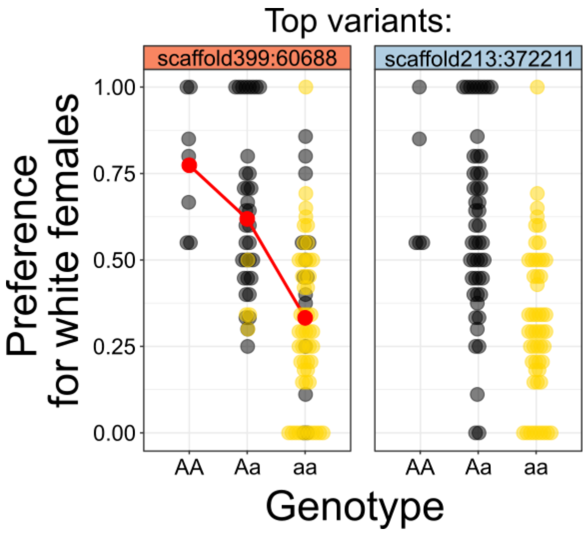


**Figure S3. Effects of allele dosage at the top preference (left) and color (right) variants on male preference (the proportion of courts directed at white females).** Each dot corresponds to a single male's lifetime preference from the original Chamberlain et al. (2009) data. Red dots indicate mean preference values for each genotype class. A/a - major and minor alleles.

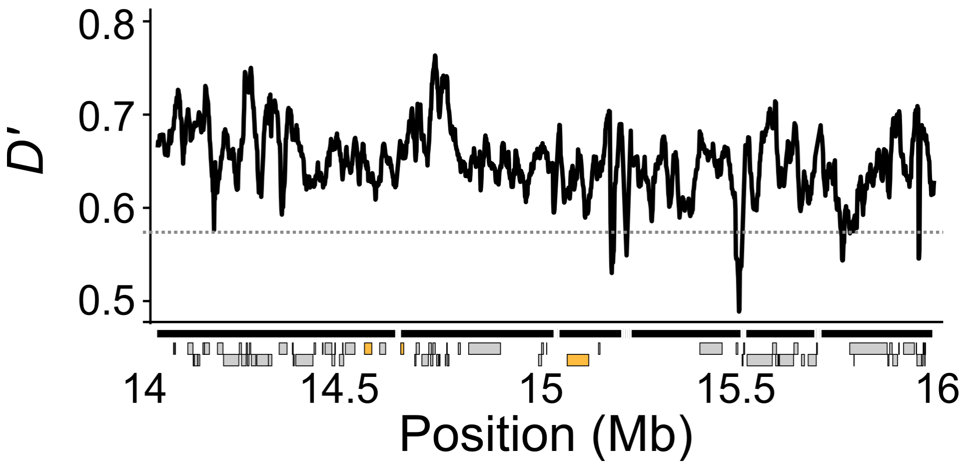


**Figure S4. *D'* across the *K* locus.** *D'* in 10 kb sliding windows (1 kb step) among the 113 sequenced *H. c. alithea* samples used in GWA. Scaffolds (black bars) and gene models (gray boxes) are shown along the *x*-axis, with (left to right) *al-1*, *al-2*, and *sens-2* filled with gold. Genome-wide average *D'* is shown as a dotted line.

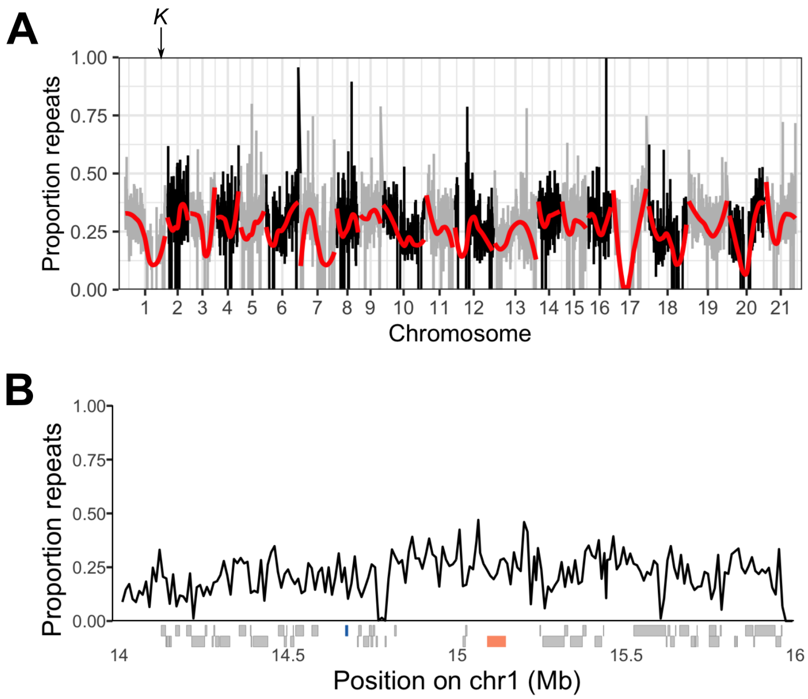


**Figure S5. The proportion of bases classified as repeats by RepeatMasker genome-wide (A) and in the *K* locus (B). A,** The proportion of masked sequence in 50 kb sliding (5 kb step) windows. Red lines represent loess fits per-chromosome. **B,** The proportion of masked sequence in 10 kb sliding (1 kb step) windows in the *K* locus. Gene models are shown as gray boxes along the *x*-axis; *al-1* and *sens-2* are highlighted in blue and gold, respectively. These results shown did not change when we limit to just putative TEs.

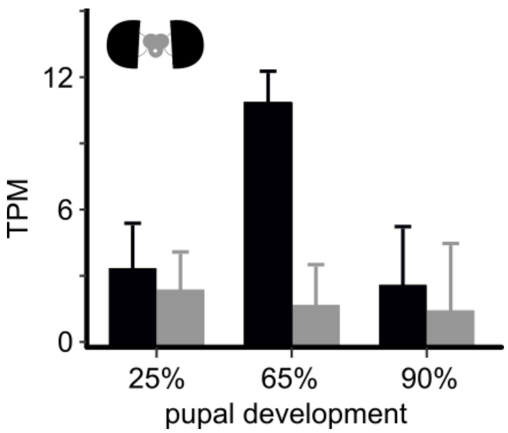

**Figure S6. Barplot showing *senseless-2* expression levels in pupal eyes (black) and central brain (gray) based on *Heliconius cydno galanthus* RNA-seq data.**

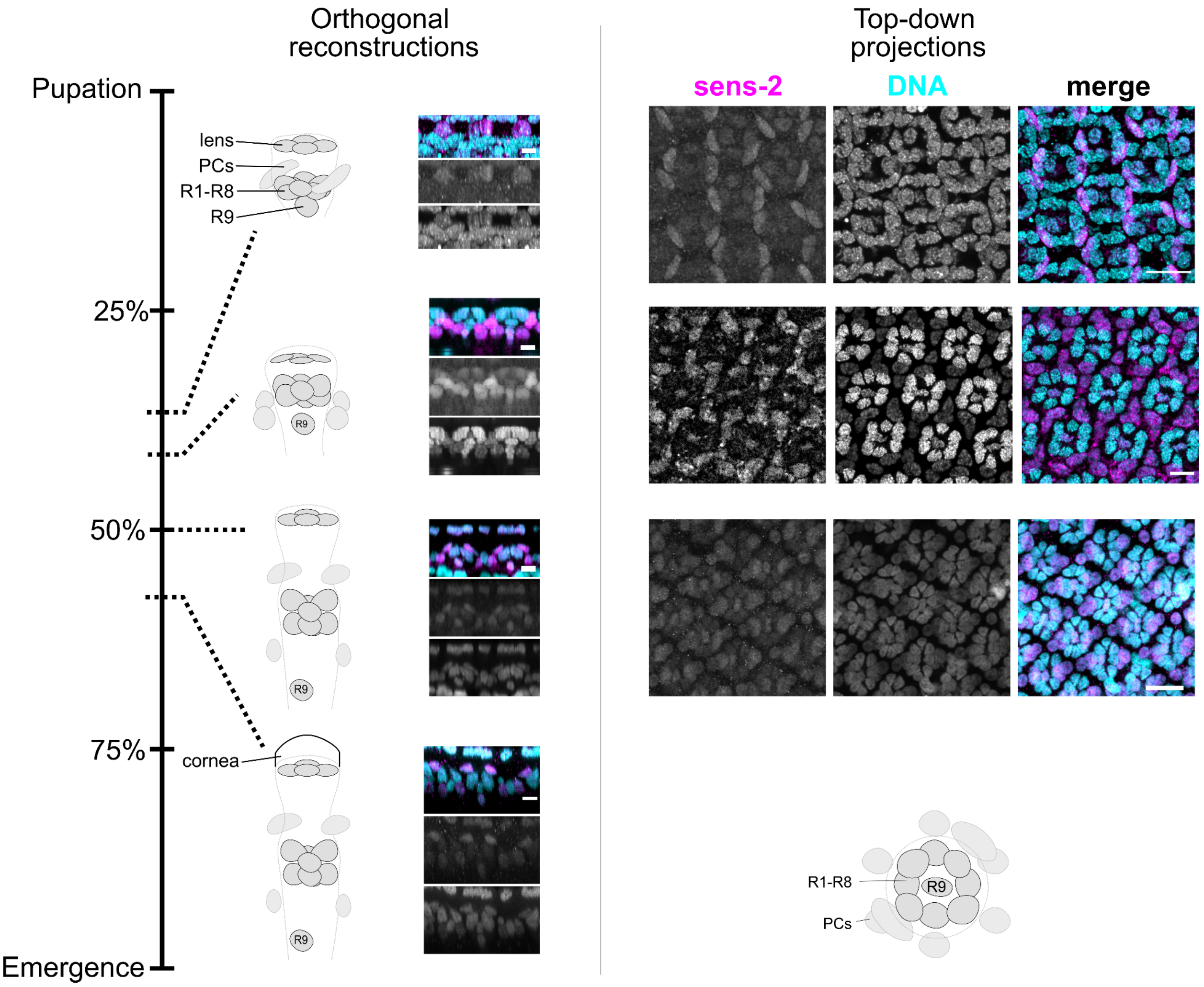


**Figure S7. Sens-2 expression patterns across mid-pupal retina development.** All individuals are yellow *Heliconius cydno alithea* males. No qualitative differences in Sens-2 staining patterns were observed between any *cydno* clade members. Scale bars: 10 um (orthogonal projections), 20 um (top-down projections).

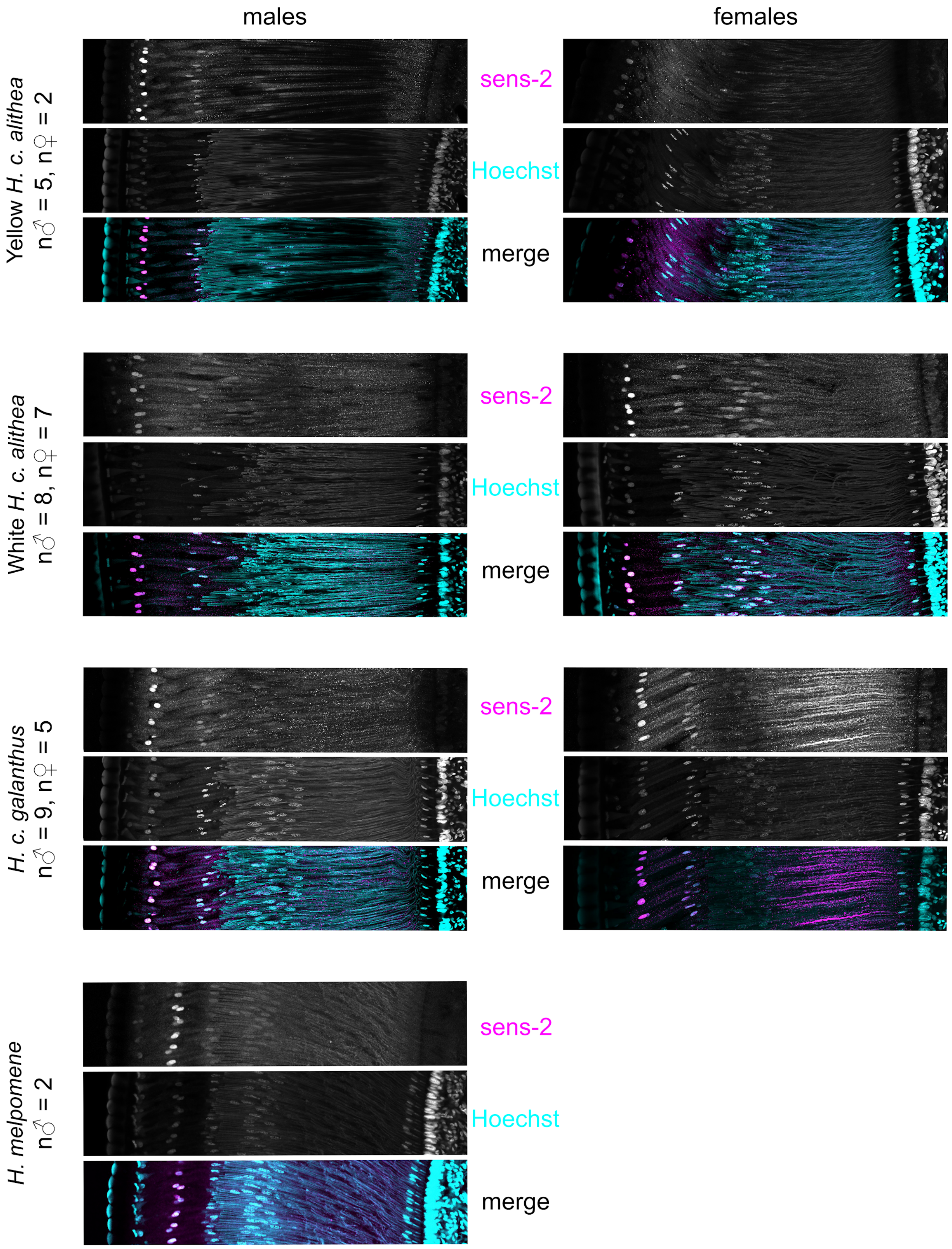


**Figure S8. Sens-2 expression in representative adult *Heliconius cydno* eyes.** All images are from adults less than 12 hours post-emergence.

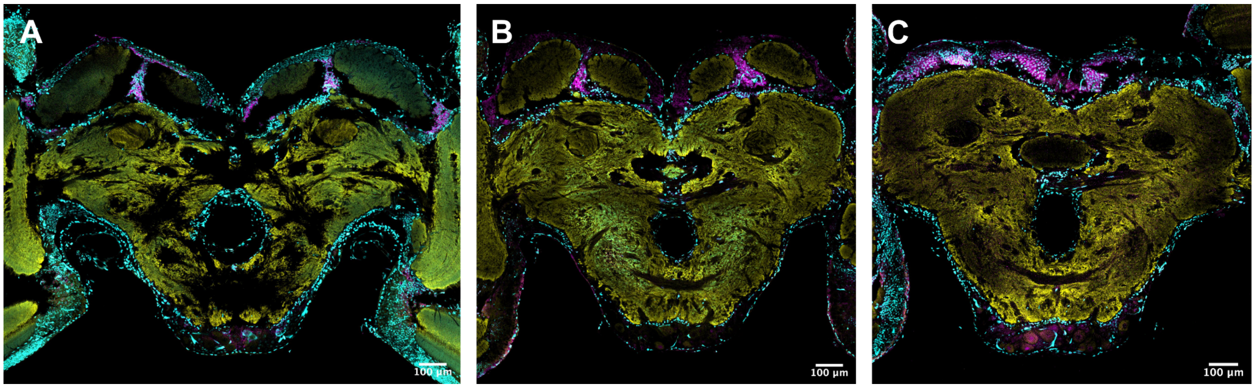


**Figure S9. Representative images of adult central brain IHC in 50 µm sections from a male white *Heliconius cydno alithea* (A), female *H. c. galanthus* (B)*,* and female yellow *H. c. alithea* (C)*.*** Cyan: DNA; magenta: Sens-2; yellow: synapsin. Mushroom body expression is consistent with weak RNAi phenotypes in Drosophila (*38*).

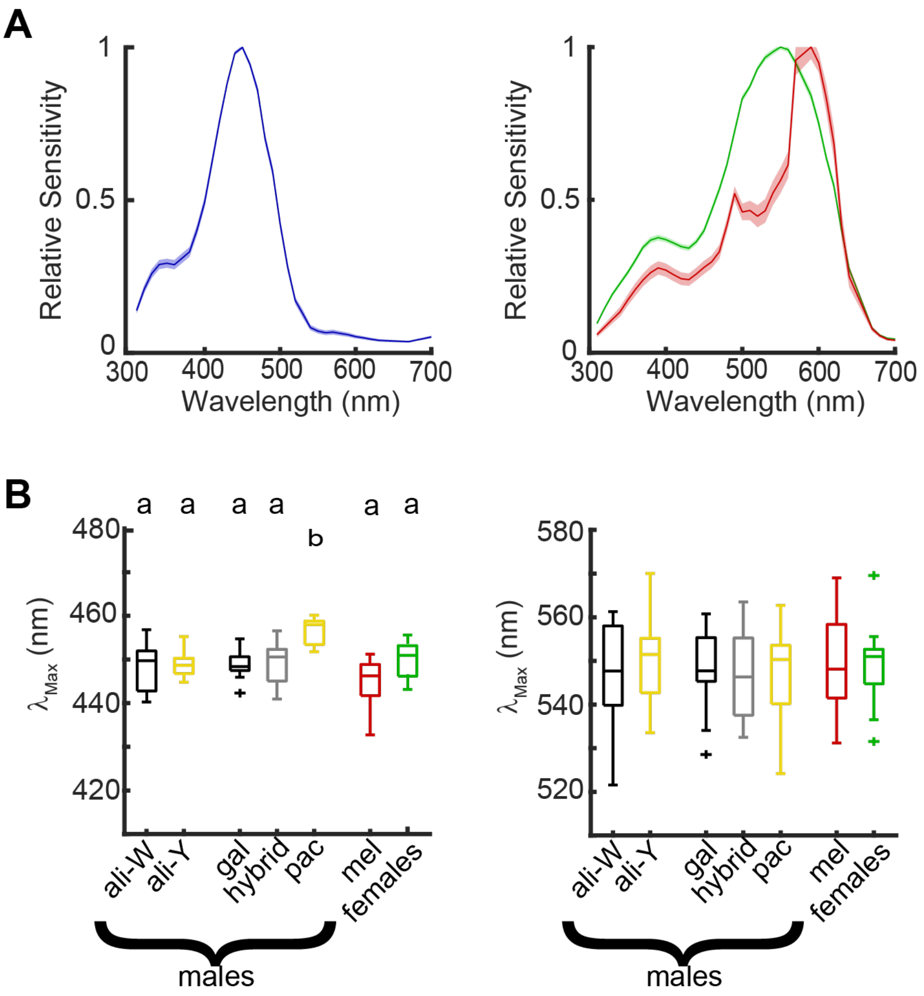


**Figure S10. Spectral tuning of blue, green, and red photoreceptors. A,** Spectral tuning curves for blue (left) and both green and red photoreceptors (right). Shading shows the standard error. **B,** Boxplots show the wavelength of peak sensitivity determined by fitting the responses of each cell with a rhodopsin template. The left shows data for blue photoreceptors (n = 21, 30, 22, 21, 5, 15, 12) and the right shows green photoreceptors (n = 27, 25, 22, 7, 11, 29, 28). Red photoreceptors are not shown due to small sample sizes (n = 2, 2, 5, 0, 1, 14, 9) and because the abnormally narrow spectral tuning that is derived from a combination of the green sensitive opsin and a red screening pigment precludes an accurate fit with the template.

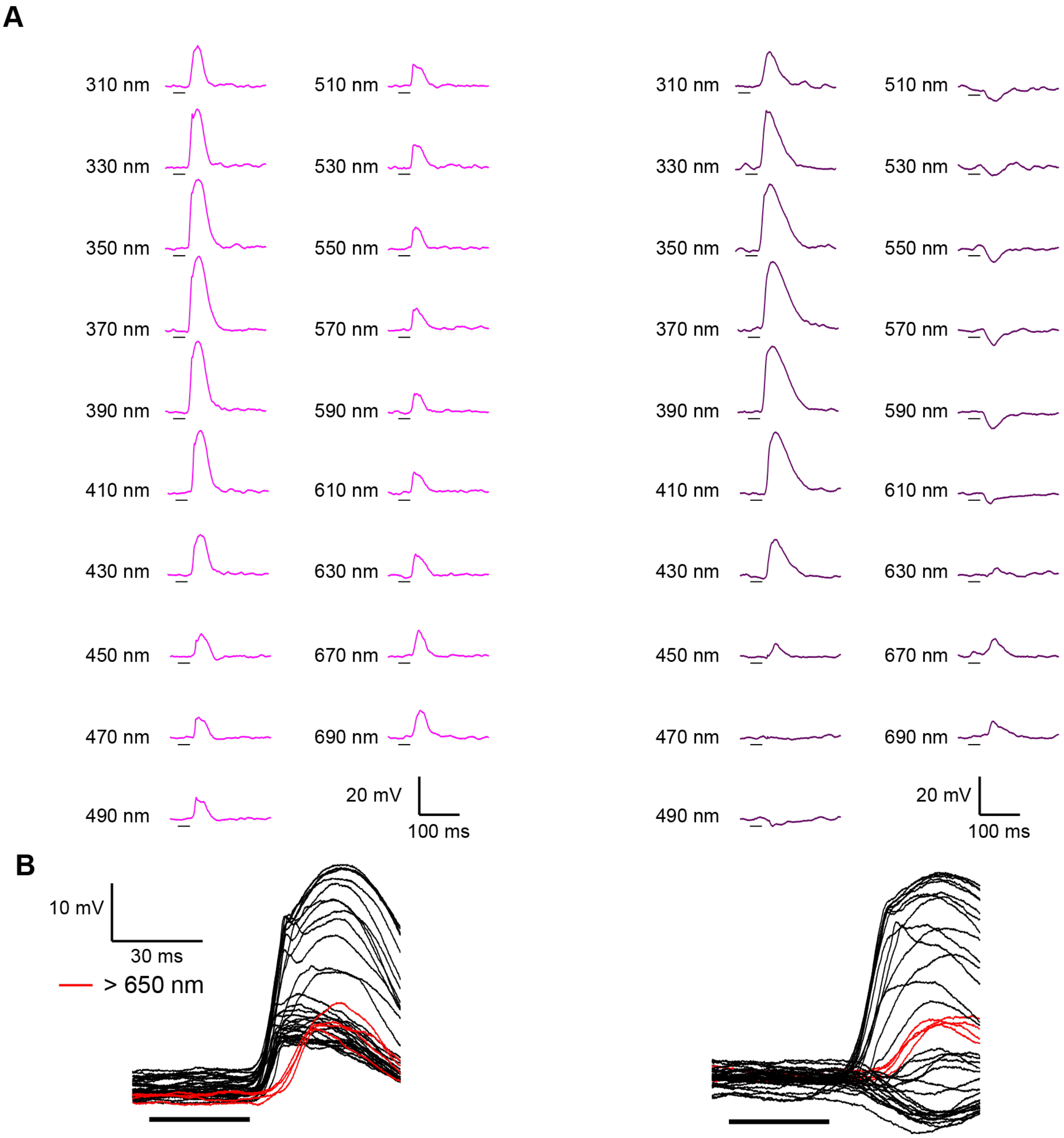


**Figure S11. Example data for UV photoreceptors with and without inhibition. A,** Data was collected using 30 ms flashes of monochromatic light in 10 nm increments. We excluded 580, 650, and 660 nm due to lamp artifacts that caused high and unstable intensities. Shown here are a subset of the responses for two UV cells from two different *H.c. alithea* males. **B,** Same data as (A), overlaid for comparison. Long wavelength stimuli are highlighted to show the delayed responses relative to both hyperpolarizing and depolarizing responses.

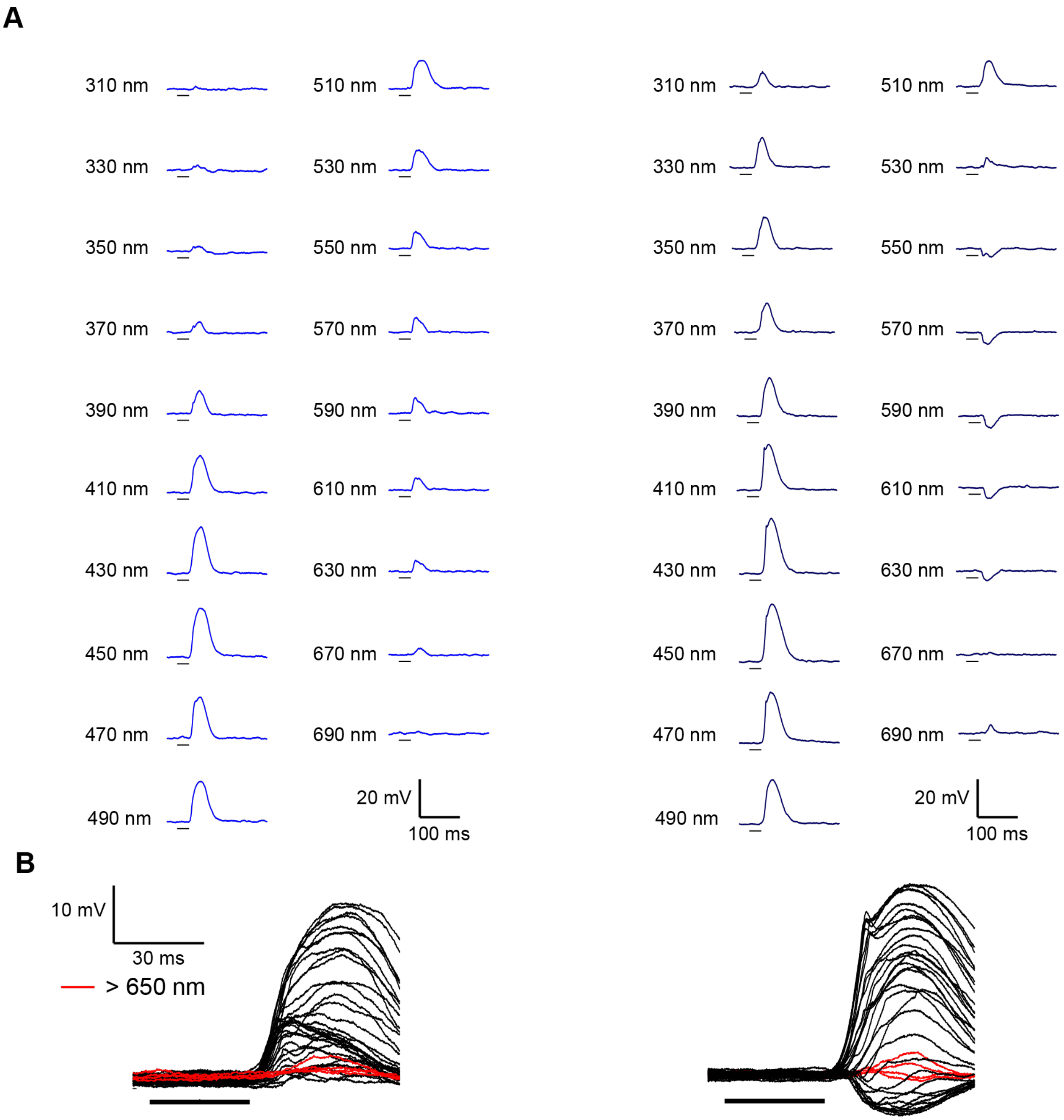


**Figure S12. Example data for blue photoreceptors with and without inhibition. A,** Traces show example data for two blue cells in two different *H.c. alithea* males. **B,** Same data as (A), overlaid for comparison. Long wavelength stimuli are highlighted to show the delayed responses relative to both hyperpolarizing and depolarizing responses.

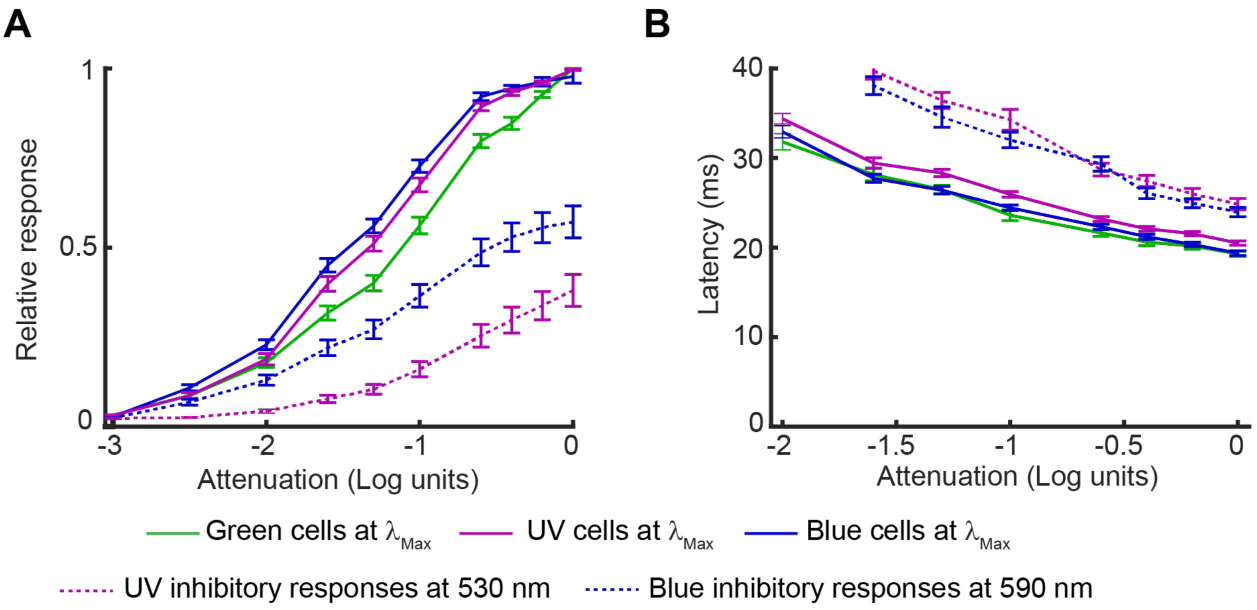


**Figure S13. Response latency data.** In addition to measuring spectral tuning, we also measured response amplitude **(A)** and latency **(B)** at different levels of light intensity. Note that panel A plots the absolute value of the response.

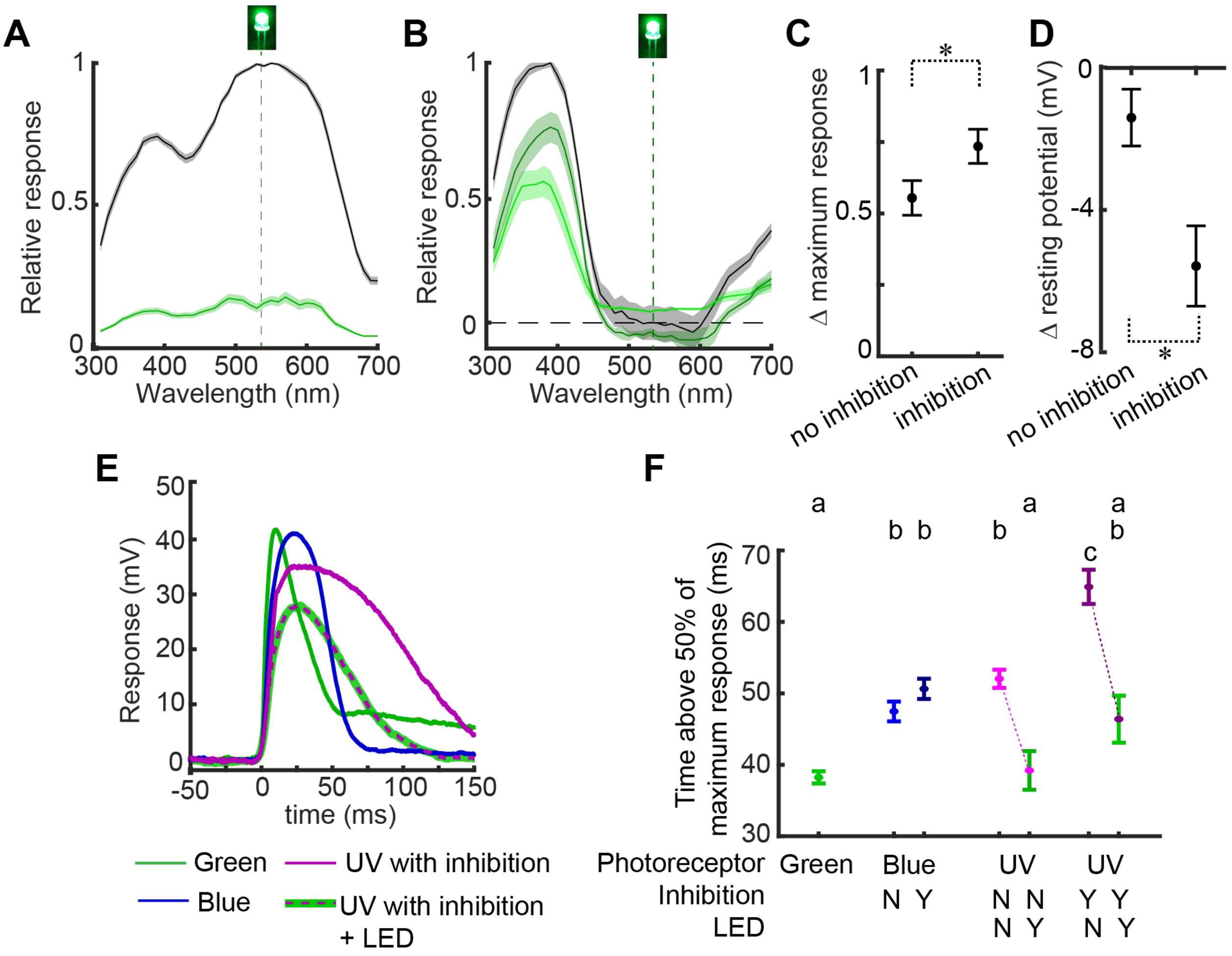


**Figure S14: UV photoreceptor responses in the presence of a 534 nm LED. A,** Response of green photoreceptors before (black) and after (green) turning on a bright green LED (dotted line). Shading shows the standard error (n=40). **B,** Response of UV photoreceptors before and after turning on the LED. Black is averaged across all UV photoreceptors before the LED, and green is separated into cells with hyperpolarizing responses (dark green, n=17) and without (light green, n=14). **C,** The addition of the LED reduced the response magnitude of UV cells without inhibition significantly more than cells with inhibition (t-test, $p < 0.01$). **D,** The LED reduced the resting potential of UV cells with inhibition (one-sample t-test, $p < 0.001$) but not UV cells without inhibition ($p = 0.10$). **E,** Raw data shows the different temporal responses of cells in different *H.c. alithea* males at the wavelength of peak sensitivity. **F,** We measured the temporal width of the response to the wavelength of peak sensitivity (time greater than 50% of the maximum voltage response). UV cells were measured both with and without the LED turned on. Letters above indicate groups significantly different from each other ($F_{6,328} = 29.92$, $p < 0.001$, Tukey's HSD). Addition of the LED long wavelength partially simulates daytime conditions, as compared to a dark adapted eye. The observed narrowing of the UV temporal response when long wavelength light is present, bringing overall PR widths in register with each other, reveals that there could be mechanisms that make PR responses more similar in naturalistic light conditions.

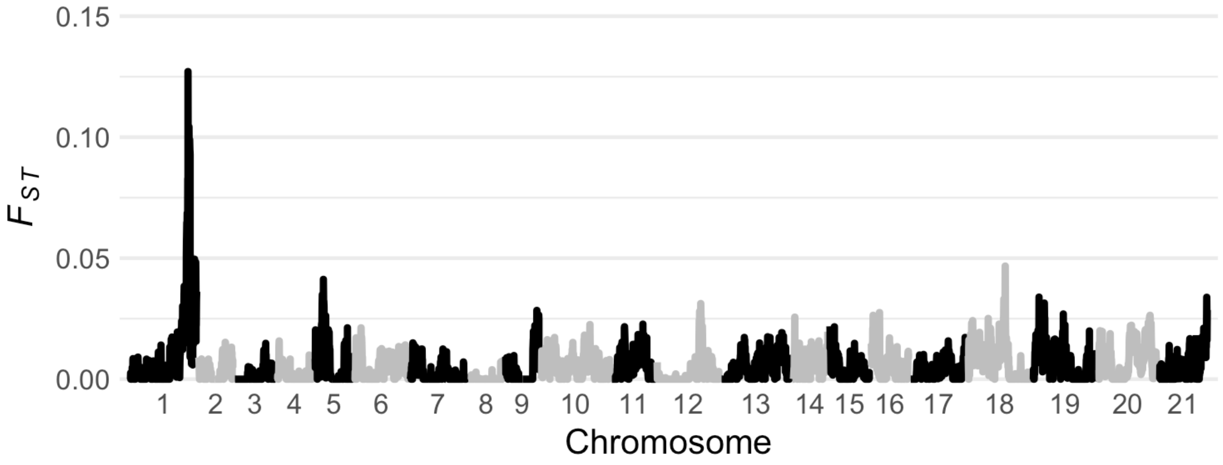


**Figure S15. Genome-wide $F_{ST}$ between yellow and white *Heliconius cydno alithea,* calculated in 50 kb sliding windows (5 kb step).** The peak on chromosome 1 corresponds to the *K* locus.

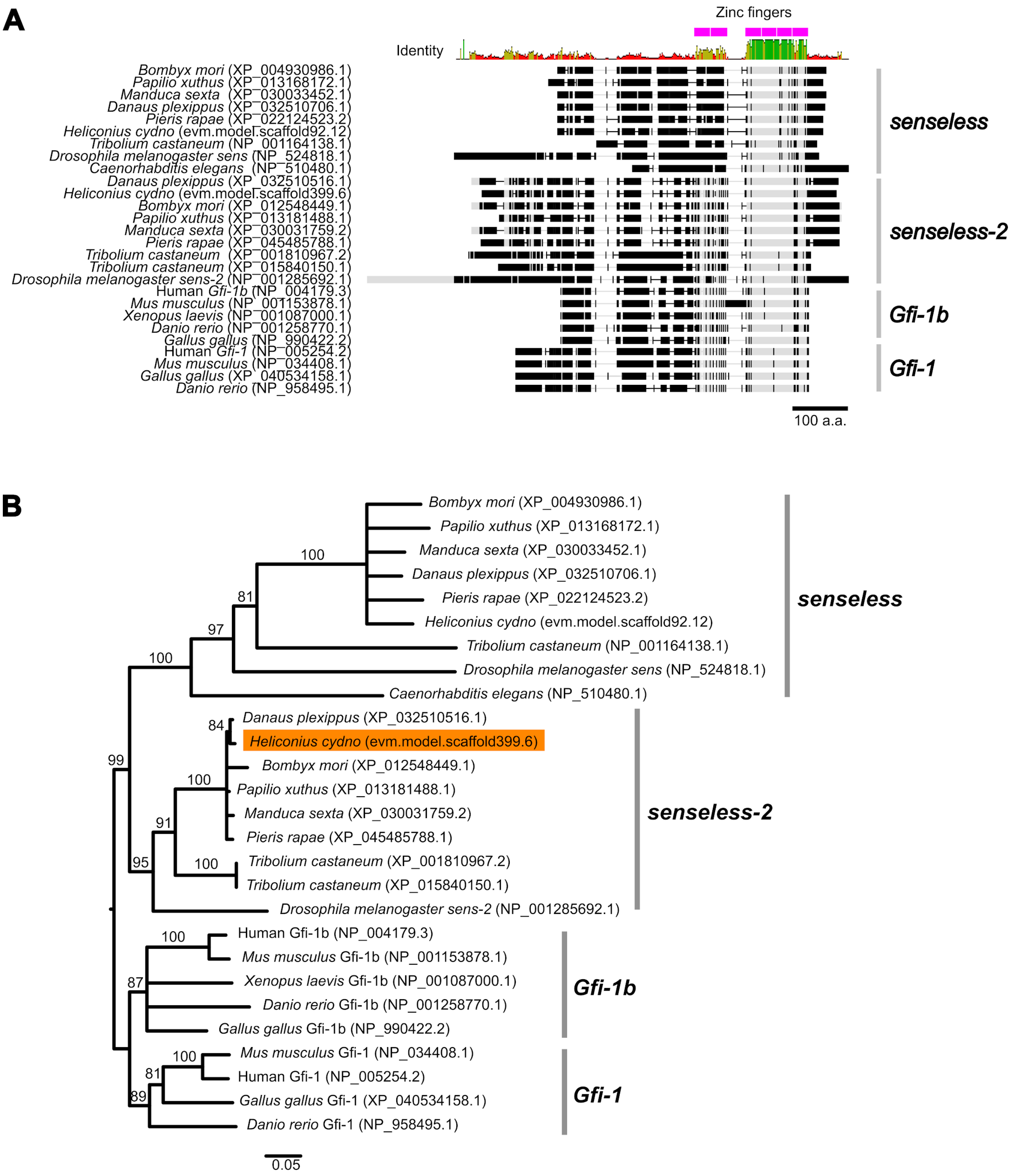


**Figure S16. Alignment and neighbor-joining tree showing *Heliconius cydno senseless-2* relationships to animal homologs.** We first used BLASTp to identify *Drosophila melanogaster* proteins most similar to the gene model scaffold399.6, the nearest gene to the top preference variants identified in the *H. c. alithea* GWA. BLASTp to the *D. melanogaster* nr database on NCBI identified *sens-2* (NP_001285692.1; E-value 6e-99) as the top hit in flies, with *senseless* (NP_524818.1; E-value 7e-54) as the second hit. A previous study showed that *Drosophila sens-2* was homologous to the vertebrate genes Gfi-1 and Gfi-1b (*39*), and we confirmed this using BLASTp against nr human proteins (top hit NP_001120687.1 Gfi-1, E-value 9e-94; second hit NP_001364233.1 Gfi-1b, E-value 6e-91). We used BLASTp to identify homologs from 11 additional well-annotated animal genomes, aligned them using MUSCLE, and constructed a neighbor-joining tree using Kimura's distance. Statistical support was assessed using 100 bootstraps. **A,** Alignment, with annotated zinc finger DNA interacting domains. Mismatches to the majority consensus sequence are shown in black. *Heliconius cydno sens-2* contains the ancestral six ZFs found in all animal orthologs, while *H. cydno sens* (scaffold92.12) contains only four ZFs. These states match those in *Drosophila melanogaster* and other invertebrates. **B,** NJ tree based on the alignment shown in **A** after stripping any sites containing gaps in any sequence.

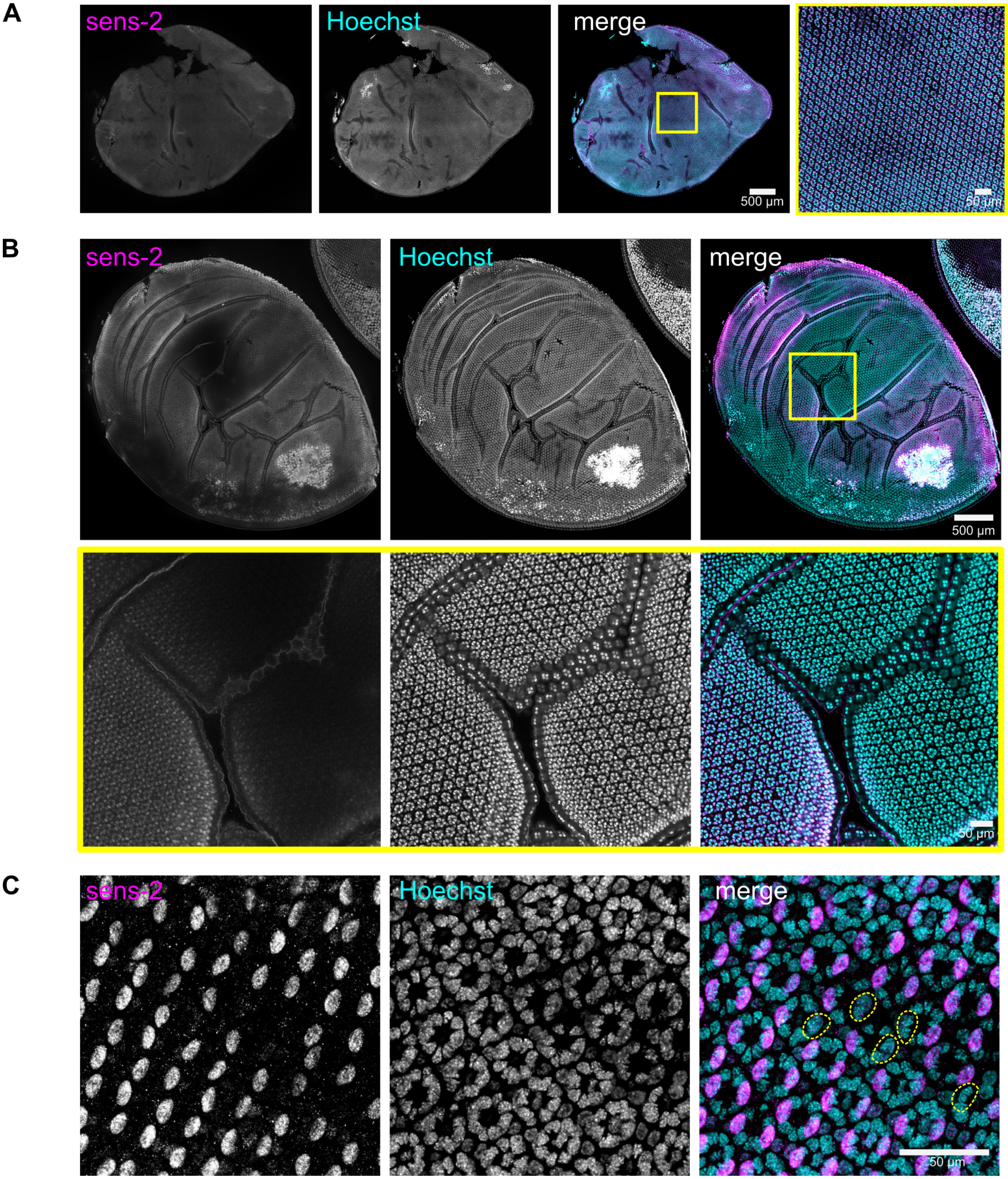


**Figure S17. Mosaic *sens-2* knockout pupal retinas. A,** Wild-type *Heliconius cydno galanthus* retina, ~50% pupal development (PD), showing uniform anti-Sens-2 staining across the retina in pigment cell nuclei (20X tile scan). **B,** Large-scale CRISPR/Cas9-induced mosaicism in a white *H. c. alithea* retina, ~55% PD. Top row: 20X tile scan. Bottom row: Zoom on yellow box. Folds are caused by rigid corneas beginning to form. **C,** Small-scale mosaicism in a ~45% PD white *H. c. alithea* retina. Compare to **A.**

**Supplementary Tables**

**Table S1.** ***Heliconius cydno alithea*** **resequencing (BioProject PRJNA802836) and phenotype information.**

| SRA Accession | Male | Forewing Color | Preference Index* | Total Courts |
|---|---|---|---|---|
| SRRXXXXXXXX | MP001 | yellow | 0.14 | 7 |
| SRRXXXXXXXX | MP002 | yellow | 0.00 | 3 |
| SRRXXXXXXXX | MP003 | yellow | 1.00 | 1 |
| SRRXXXXXXXX | MP004 | yellow | 0.63 | 8 |
| SRRXXXXXXXX | MP005 | yellow | 0.45 | 11 |
| SRRXXXXXXXX | MP006 | yellow | 0.50 | 30 |
| SRRXXXXXXXX | MP008 | white | 0.40 | 43 |
| SRRXXXXXXXX | MP009 | white | 1.00 | 1 |
| SRRXXXXXXXX | MP010 | white | 0.75 | 4 |
| SRRXXXXXXXX | MP012 | yellow | 0.00 | 1 |
| SRRXXXXXXXX | MP013 | white | 0.71 | 14 |
| SRRXXXXXXXX | MP016 | yellow | 0.00 | 1 |
| SRRXXXXXXXX | MP019 | white | 0.00 | 1 |
| SRRXXXXXXXX | MP020 | white | 1.00 | 1 |
| SRRXXXXXXXX | MP022 | yellow | 0.50 | 2 |
| SRRXXXXXXXX | MP025 | white | 1.00 | 1 |
| SRRXXXXXXXX | MP026 | yellow | 0.30 | 23 |
| SRRXXXXXXXX | MP027 | yellow | 0.33 | 3 |
| SRRXXXXXXXX | MP028 | yellow | 0.00 | 1 |
| SRRXXXXXXXX | MP029 | yellow | 0.00 | 1 |
| SRRXXXXXXXX | MP031 | white | 0.50 | 12 |
| SRRXXXXXXXX | MP033 | yellow | 0.30 | 29 |
| SRRXXXXXXXX | MP035 | white | 0.55 | 30 |
| SRRXXXXXXXX | MP037 | yellow | 0.20 | 32 |
| SRRXXXXXXXX | MP038 | white | 0.80 | 22 |
| SRRXXXXXXXX | MP039 | white | 0.70 | 23 |
| SRRXXXXXXXX | MP040 | yellow | 0.60 | 29 |
| SRRXXXXXXXX | MP041 | yellow | 0.65 | 23 |
| SRRXXXXXXXX | MP042 | white | 0.30 | 24 |
| SRRXXXXXXXX | MP043 | yellow | 0.25 | 20 |
| SRRXXXXXXXX | MP044 | yellow | 0.55 | 22 |
| SRRXXXXXXXX | MP045 | white | 1.00 | 2 |
| SRRXXXXXXXX | MP046 | yellow | 0.35 | 23 |
| SRRXXXXXXXX | MP048 | yellow | 0.35 | 21 |
| SRRXXXXXXXX | MP049 | yellow | 0.15 | 21 |
| SRRXXXXXXXX | MP050 | white | 0.85 | 26 |
| SRRXXXXXXXX | MP051 | yellow | 0.50 | 29 |
| SRRXXXXXXXX | MP052 | yellow | 0.55 | 22 |
| SRRXXXXXXXX | MP053 | yellow | 0.35 | 27 |
| SRRXXXXXXXX | MP054 | white | 0.45 | 26 |
| SRRXXXXXXXX | MP058 | yellow | 0.00 | 1 |
| SRRXXXXXXXX | MP059 | white | 0.67 | 6 |
| SRRXXXXXXXX | MP064 | yellow | 0.25 | 23 |

**Table S1. Continued**

| SRA Accession | Male | Forewing Color | Preference Index* | Total Courts |
|---|---|---|---|---|
| SRRXXXXXXXX | MP067 | white | 0.55 | 24 |
| SRRXXXXXXXX | MP068 | white | 0.64 | 14 |
| SRRXXXXXXXX | MP069 | white | 0.55 | 22 |
| SRRXXXXXXXX | MP070 | white | 0.80 | 23 |
| SRRXXXXXXXX | MP071 | white | 0.25 | 23 |
| SRRXXXXXXXX | MP073 | yellow | 0.00 | 1 |
| SRRXXXXXXXX | MP075 | white | 0.60 | 24 |
| SRRXXXXXXXX | MP076 | white | 0.70 | 20 |
| SRRXXXXXXXX | MP078 | white | 0.44 | 9 |
| SRRXXXXXXXX | MP080 | white | 0.25 | 20 |
| SRRXXXXXXXX | MP081 | white | 0.75 | 20 |
| SRRXXXXXXXX | MP082 | yellow | 0.21 | 19 |
| SRRXXXXXXXX | MP083 | yellow | 0.50 | 24 |
| SRRXXXXXXXX | MP085 | white | 0.55 | 22 |
| SRRXXXXXXXX | MP086 | yellow | 0.25 | 25 |
| SRRXXXXXXXX | MP087 | white | 0.60 | 23 |
| SRRXXXXXXXX | MP089 | yellow | 0.30 | 25 |
| SRRXXXXXXXX | MP090 | white | 0.11 | 9 |
| SRRXXXXXXXX | MP091 | white | 0.40 | 23 |
| SRRXXXXXXXX | MP093 | white | 0.40 | 22 |
| SRRXXXXXXXX | MP094 | yellow | 0.25 | 8 |
| SRRXXXXXXXX | MP095 | white | 0.86 | 7 |
| SRRXXXXXXXX | MP096 | yellow | 0.33 | 3 |
| SRRXXXXXXXX | MP097 | white | 0.50 | 6 |
| SRRXXXXXXXX | MP098 | white | 0.50 | 8 |
| SRRXXXXXXXX | MP099 | white | 0.55 | 21 |
| SRRXXXXXXXX | MP101 | yellow | 0.50 | 2 |
| SRRXXXXXXXX | MP102 | white | 0.55 | 22 |
| SRRXXXXXXXX | MP103 | white | 0.45 | 21 |
| SRRXXXXXXXX | MP107 | white | 0.45 | 21 |
| SRRXXXXXXXX | MP108 | white | 0.55 | 22 |
| SRRXXXXXXXX | MP109 | yellow | 0.00 | 1 |
| SRRXXXXXXXX | MP110 | yellow | 0.45 | 35 |
| SRRXXXXXXXX | MP111 | yellow | 0.45 | 21 |
| SRRXXXXXXXX | MP112 | white | 0.67 | 18 |
| SRRXXXXXXXX | MP113 | yellow | 0.41 | 17 |
| SRRXXXXXXXX | MP114 | white | 0.38 | 16 |
| SRRXXXXXXXX | MP115 | yellow | 0.19 | 16 |
| SRRXXXXXXXX | MP116 | white | 0.64 | 14 |
| SRRXXXXXXXX | MP117 | yellow | 0.35 | 24 |
| SRRXXXXXXXX | MP118 | white | 0.45 | 20 |
| SRRXXXXXXXX | MP120 | yellow | 0.30 | 20 |
| SRRXXXXXXXX | MP121 | yellow | 0.60 | 22 |
| SRRXXXXXXXX | MP122 | white | 0.50 | 10 |
| SRRXXXXXXXX | MP123 | yellow | 0.50 | 6 |
| SRRXXXXXXXX | MP124 | yellow | 0.69 | 13 |

**Table S1. Continued**

| SRA Accession | Male | Forewing Color | Preference Index* | Total Courts |
|---|---|---|---|---|
| SRRXXXXXXXX | MP125 | white | 0.75 | 4 |
| SRRXXXXXXXX | MP127 | yellow | 0.25 | 4 |
| SRRXXXXXXXX | MP128 | white | 0.50 | 8 |
| SRRXXXXXXXX | MP131 | yellow | 0.20 | 5 |
| SRRXXXXXXXX | MP132 | yellow | 0.50 | 2 |
| SRRXXXXXXXX | MP133 | white | 1.00 | 2 |
| SRRXXXXXXXX | MP134 | yellow | 0.18 | 17 |
| SRRXXXXXXXX | MP136 | yellow | 0.43 | 14 |
| SRRXXXXXXXX | MP137 | white | 1.00 | 1 |
| SRRXXXXXXXX | MP139 | white | 1.00 | 1 |
| SRRXXXXXXXX | MP140 | white | 0.33 | 6 |
| SRRXXXXXXXX | MP142 | yellow | 0.14 | 7 |
| SRRXXXXXXXX | MP143 | yellow | 0.00 | 4 |
| SRRXXXXXXXX | MP145 | yellow | 0.29 | 7 |
| SRRXXXXXXXX | MP147 | yellow | 0.00 | 1 |
| SRRXXXXXXXX | MP153 | yellow | 0.20 | 5 |
| SRRXXXXXXXX | MP155 | white | 1.00 | 1 |
| SRRXXXXXXXX | MP156 | yellow | 0.30 | 21 |
| SRRXXXXXXXX | MP162 | white | 0.00 | 1 |
| SRRXXXXXXXX | MP163 | white | 0.33 | 3 |
| SRRXXXXXXXX | MP165 | white | 0.80 | 5 |
| SRRXXXXXXXX | MP167 | yellow | 0.00 | 2 |
| SRRXXXXXXXX | MP169 | white | 1.00 | 2 |
| SRRXXXXXXXX | MP171 | white | 1.00 | 1 |

*Preference Index = # courts toward white females / total courts (up to 20 courts - affects 36 males).

**Table S2. Gene models shown in Fig. 1E. See Supplementary Methods for details on genome assembly and annotation.**

| | | | BLASTp to NCBI nr database | | | | | |
|---|---|---|---|---|---|---|---|---|
| Gene (evm.TU) | Name* | Strand | Accession | Description | Subject Species | Query Cov. | E-value | %ID |
| scaffold57.33 | - | + | XP_038222035.1 | uncharacterized protein | Zerene cesonia | 80% | 1e-99 | 88.89% |
| scaffold57.32 | *ttc14* | + | XP_030041110.1 | tetratricopeptide repeat protein 14 homolog | Manduca sexta | 100% | 0 | 85.87% |
| scaffold57.31 | *mpdu1* | - | XP_028175378.1 | mannose-P-dolichol utilization defect 1 protein homolog | Ostrinia furnacalis | 94% | 1e-134 | 80.26% |
| scaffold57.30 | - | - | CAG9567441.1 | unnamed protein product | Danaus chrysippus | 100% | 3e-107 | 94.47% |
| scaffold57.29 | *ttll8* | - | XP_026487438.1 | tubulin glycylase 3A-like | Vanessa tameamea | 99% | 0 | 63.59% |
| scaffold57.26 | *stk24* | + | XP_026487821.1 | serine/threonine-protein kinase 26 | - | 99% | 0 | 91.42% |
| scaffold57.25 | - | - | XP_026487397.1 | integrin beta pat-3-like | - | 88% | 3e-166 | 59.66% |
| scaffold57.23 | - | - | XP_026487395.1 | integrin beta-PS-like | Vanessa tameamea | 99% | 0 | 57.01% |
| scaffold57.22 | - | + | XP_026487576.1 | integrin beta pat-3-like | Vanessa tameamea | 98% | 0 | 57.71% |
| scaffold57.21 | - | - | XP_032530016.1 | putative leucine-rich repeat-containing protein | Danaus plexippus | 99% | 0 | 46.32% |
| scaffold57.20 | - | + | XP_026487535.1 | uncharacterized protein | Vanessa tameamea | 97% | 7e-116 | 50.41% |
| scaffold57.19 | - | - | XP_026487533.1 | lipase 3-like | Vanessa tameamea | 100% | 0 | 75.49% |
| scaffold57.18 | - | + | XP_026487436.1 | zinc finger protein 131-like, bric-a-brac, longitudinals lacking | Vanessa tameamea | 100% | 0 | 83.25% |
| scaffold57.16 | *ninaC* | - | QDR50934.1 | neither inactivation nor afterpotential C | Heliconius melpomene | 100% | 4e-67 | 100.00% |
| scaffold57.15 | *Wnt10a* | - | XP_026487660.1 | protein Wnt-10a | Vanessa tameamea | 96% | 0 | 87.63% |
| scaffold57.14 | *Wnt6* | + | XP_032510632.1 | protein Wnt-6 | Danaus plexippus | 100% | 0 | 92.61% |
| scaffold57.13 | - | - | XP_032510631.1 | protein PF14_0175-like | Danaus plexippus | 59% | 2e-93 | 54.92% |
| scaffold57.12 | *Wnt1/wg* | - | XP_032530256.1 | protein Wnt-1 | Danaus plexippus | 100% | 0 | 94.64% |
| scaffold57.11 | *Wnt4* | - | XP_034825530.1 | protein Wnt-4 | Maniola hyperantus | 100% | 0 | 78.59% |
| scaffold57.8 | *SLC4A7* | + | XP_026487485.1 | sodium bicarbonate cotransporter 3 | Vanessa tameamea | 100% | 0 | 94.23% |

| | | | | | | | | |
|---|---|---|---|---|---|---|---|---|
| scaffold57.7 | *gfm1* | - | XP_026487492.1 | elongation factor G, mitochondrial | Vanessa tameamea | 100% | 0 | 90.20% |
| novel_TU_984_60854d56 | *PRKACA* | - | XP_032510666.1 | cAMP-dependent protein kinase catalytic subunit alpha-like | Danaus plexippus | 100% | 9e-69 | 81.67% |
| temp_TU_280.1.608a89df | - | - | XP_026487509.1 | uncharacterized protein | Vanessa tameamea | 96% | 5e-100 | 65.70% |
| scaffold57.6 | *lrch3* | + | XP_026487491.1 | leucine-rich repeat and calponin homology domain-containing protein 3 | Vanessa tameamea | 97% | 0 | 88.52% |
| scaffold57.4 | *aristaless-2* | + | ADA70355.1 | paired-like family homeodomain transcription factor | Heliconius erato | 100% | 0 | 99.61% |
| scaffold57.2 | - | + | XP_034824464.1 | homeobox protein aristaless-like | Maniola hyperantus | 39% | 2e-18 | 71.83% |
| scaffold213.23 | *aristaless-1* | + | ADA70356.1 | paired-like family homeodomain transcription factor | Heliconius erato | 100% | 0 | 99.25% |
| scaffold213.19 | *eIF3h* | - | XP_026487502.1 | eukaryotic translation initiation factor 3 subunit H | Vanessa tameamea | 100% | 0 | 97.03% |
| scaffold213.18 | *alsin-2* | + | XP_026487482.1 | alsin | Vanessa tameamea | 100% | 0 | 86.66% |
| scaffold213.17 | *ctdspl* | - | XP_026757277.1 | carboxy-terminal domain RNA polymerase II polypeptide A small phosphatase 1 | Galleria mellonella | 100% | 0 | 97.60% |
| scaffold213.16 | *rngtt* | + | XP_034827857.1 | mRNA-capping enzyme | Maniola hyperantus | 99% | 0 | 66.61% |
| scaffold213.15 | *lactb2* | - | XP_041969043.1 | beta-lactamase-like protein 2 homolog | Aricia agestis | 100% | 4e-180 | 80.55% |
| scaffold213.14 | *ndufv1* | + | XP_026487495.1 | NADH dehydrogenase [ubiquinone] flavoprotein 1, mitochondrial | Vanessa tameamea | 100% | 0 | 96.01% |
| scaffold213.12 | *trb2* | - | XP_026487500.1 | tribbles homolog 2-like | Vanessa tameamea | 100% | 0 | 97.32% |
| scaffold213.9 | *tFIIB* | + | XP_004929188.2 | transcription initiation factor IIB | Bombyx mori | 97% | 0 | 97.21% |
| scaffold213.4 | *HEATR1* | - | XP_026488769.1 | HEAT repeat-containing protein 1 | Vanessa tameamea | 100% | 0 | 77.10% |
| scaffold213.3 | *ppt2* | + | XP_032529545.1 | lysosomal thioesterase PPT2 homolog | Danaus plexippus | 100% | 0 | 87.37% |
| scaffold213.2 | - | + | XP_026488781.1 | uncharacterized protein | Vanessa tameamea | 100% | 7e-71 | 84.55% |
| scaffold399.6 | *senseless-2* | - | XP_023942153.1 | zinc finger protein Gfi-1 | Bicyclus anynana | 99% | 0 | 90.68% |

| | | | | | | | | |
|---|---|---|---|---|---|---|---|---|
| scaffold399.5 | *rpusd2* | + | XP_034841050.1 | RNA pseudouridylate synthase domain-containing protein 2-like | Maniola hyperantus | 53% | 1e-34 | 66.33% |
| scaffold399.1 | - | - | CAG4954309.1 | unnamed protein product | Parnassius apollo | 32% | 5e-07 | 58.70% |
| scaffold376.1 | *La* | + | XP_026488796.1 | la protein homolog | Vanessa tameamea | 98% | 2e-97 | 80.73% |
| scaffold376.2 | *SAAL1* | + | XP_023942364.1 | uncharacterized protein | Bicyclus anynana | 100% | 2e-46 | 72.07% |
| scaffold376.3 | *sky* | - | XP_026488789.1 | TBC1 domain family member 24 | Vanessa tameamea | 100% | 0 | 98.30% |
| scaffold376.4 | *mroh1* | + | XP_026488783.1 | maestro heat-like repeat-containing protein family member 1 | Vanessa tameamea | 98% | 0 | 85.81% |
| scaffold376.5 | - | - | XP_031763815.1 | uncharacterized protein | Galleria mellonella | 96% | 1e-70 | 56.63% |
| scaffold376.6 | - | - | XP_026488770.1 | uncharacterized protein | Vanessa tameamea | 98% | 2e-71 | 54.95% |
| scaffold376.7 | *eepd1* | - | XP_026488793.1 | endonuclease/exonuclease/ phosphatase family domain-containing protein 1-like | Vanessa tameamea | 100% | 0 | 80.54% |
| scaffold376.8 | *DYNC 1LI1* | - | XP_026488794.1 | cytoplasmic dynein 1 light intermediate chain 2 | Vanessa tameamea | 100% | 0 | 95.26% |
| scaffold376.9 | - | + | XP_039764575.1 | coiled-coil and C2 domain-containing protein 1-like | Pararge aegeria | 100% | 0 | 67.92% |
| scaffold376.12 | *KLF11* | - | XP_035446396.1 | Krueppel-like factor 10 | Spodoptera frugiperda | 97% | 1e-169 | 81.93% |
| scaffold894.1 | - | + | XP_026486895.1 | putative inactive cysteine synthase 2 | Vanessa tameamea | 98% | 0 | 85.00% |
| scaffold160.25 | *u-shaped* | + | XP_026490476.1 | zinc finger protein ush | Vanessa tameamea | 100% | 0 | 89.70% |
| scaffold160.24 | *SLC7A 6OS* | - | XP_026490510.1 | probable RNA polymerase II nuclear localization protein SLC7A6OS | Vanessa tameamea | 100% | 1e-155 | 75.25% |
| scaffold160.23 | *MED15* | + | XP_034834684.1 | mediator of RNA polymerase II transcription subunit 15 | Maniola hyperantus | 100% | 0 | 87.09% |
| scaffold160.22 | *TRIM4 5* | - | XP_039764764.1 | tripartite motif-containing protein 45 | Pararge aegeria | 98% | 0 | 71.07% |
| scaffold160.21 | *HS2ST 1* | + | XP_026490505.1 | heparin sulfate O-sulfotransferase | Vanessa tameamea | 81% | 3e-72 | 86.89% |
| scaffold160.20 | *Adk1-1* | + | XP_026490523.1 | uncharacterized protein | Vanessa tameamea | 100% | 7e-137 | 94.23% |

| | | | | | | | | |
|---|---|---|---|---|---|---|---|---|
| scaffold160.18 | *TGIF1* | - | XP_023941033.1 | homeobox protein PKNOX1-like | Bicyclus anynana | 99% | 8e-77 | 60.96% |
| scaffold160.16 | *Adk1-2* | - | XP_026490497.1 | adenylate kinase isoenzyme 1-like | Vanessa tameamea | 98% | 1e-108 | 88.76% |
| scaffold160.15 | - | + | XP_026490496.1 | uncharacterized protein | Vanessa tameamea | 96% | 5e-99 | 64.32% |
| scaffold160.14 | *CIB1* | + | XP_034835125.1 | calcium and integrin-binding protein 1-like | Maniola hyperantus | 100% | 1e-130 | 98.38% |
| scaffold160.13 | *OSTC* | - | XP_026490511.1 | oligosaccharyltransferase complex subunit ostc-A | Vanessa tameamea | 100% | 3e-102 | 97.99% |
| scaffold160.12 | *SUCO* | + | XP_026490442.1 | SUN domain-containing ossification factor | Vanessa tameamea | 100% | 0 | 79.88% |
| scaffold160.10 | - | + | XP_026500685.1 | uncharacterized protein | Vanessa tameamea | 98% | 7e-48 | 53.12% |
| scaffold160.11 | *METTL13* | - | XP_026490448.1 | methyltransferase-like protein 13 | Vanessa tameamea | 100% | 0 | 81.61% |
| scaffold160.9 | *eIF2ak1* | + | XP_039764833.1 | eukaryotic translation initiation factor 2-alpha kinase 1-like | Pararge aegeria | 99% | 0 | 78.79% |
| scaffold160.8 | *DYRK4* | - | XP_026490447.1 | basic-leucine zipper transcription factor A | Vanessa tameamea | 100% | 0 | 83.66% |
| scaffold160.7 | *DYRK2* | - | XP_034833183.1 | dual specificity tyrosine-phosphorylation-regulated kinase 2 | Maniola hyperantus | 94% | 0 | 96.57% |
| scaffold160.6 | *DOCK6* | + | XP_034828136.1 | dedicator of cytokinesis protein 7 | Maniola hyperantus | 100% | 0 | 90.30% |
| scaffold160.5 | *cbx1* | - | XP_021197108.1 | chromobox protein homolog 5-like | Helicoverpa armigera | 100% | 2e-102 | 81.35% |
| scaffold160.4 | *nfat5* | + | XP_026490470.1 | uncharacterized protein | Vanessa tameamea | 100% | 0 | 79.19% |
| scaffold160.3 | *preb* | - | XP_026490475.1 | prolactin regulatory element-binding protein | Vanessa tameamea | 100% | 0 | 87.89% |
| scaffold160.2 | - | - | XP_026490474.1 | uncharacterized protein | Vanessa tameamea | 100% | 0 | 88.34% |
| scaffold160.1 | *OCRL* | + | XP_026490473.1 | type II inositol 1,4,5-trisphosphate 5-phosphatase | Vanessa tameamea | 100% | 0 | 91.67% |
| scaffold29A.3 | *paramyosin* | + | XP_026490483.1 | paramyosin | Vanessa tameamea | 50% | 0 | 96.72% |

*Based on BLASTp results and eggNOG annotations

**Table S3.** ***Heliconius cydno galanthus*** **genome sequencing data (PRJNA802828).**

| SRA Accession | Insert Size | Format | Read Pairs (M) |
|---|---|---|---|
| SRR17923070 | 500 bp | paired-end | 66.9 |
| SRR17923071 | 500 bp | paired-end | 14.2 |
| SRR17923072 | 500 bp | paired-end | 25.4 |
| SRR17923073 | 500 bp | paired-end | 98.3 |
| SRR17923074 | 6 kb | mate pair | 19.6 |
| SRR17923075 | 2 kb | mate pair | 35.9 |
| SRR17923076 | 15 kb | mate pair | 19.6 |
| SRR17923077 | 250 bp | paired-end | 183.0 |

**Table S4. Evidence weights used in EVM-based annotation of the *H. c. galanthus* genome.**

| Evidence | Type | Weight |
|---|---|---|
| GlimmerHMM | ABINITIO_PREDICTION | 1 |
| Genemark-ET | ABINITIO_PREDICTION | 1 |
| Augustus | ABINITIO_PREDICTION | 1 |
| High-Quality Augustus | OTHER_PREDICTION | 4 |
| TransDecoder Assemblies | OTHER_PREDICTION | 10 |
| Exonerate | PROTEIN | 4 |
| PASA Assemblies | TRANSCRIPT | 5 |

**Table S5. Oligonucleotides used in this study.**

| Name | Sequence | Type | Note |
|---|---|---|---|
| sens2_HRM_1F | AATAAGTGCTATGCCGCCGA | screening primer | - |
| sens2_HRM_2F | ACTGATCGCACGTCGGTTTA | screening primer | - |
| sens2_HRM_1R | CTGAAAGCGCTATTGCTCGG | screening primer | - |
| sens2_HRM_2R | AGCTTACCTTTCGGTGTGACA | screening primer | - |
| QPLH2711.AD | AGGAGCAGTTCTCCAAGAGA | sgRNA target | 48 bp after start (+) |
| QPLH2711.AE | CTCCAAGAGAAGGATCTCCA | sgRNA target | 58 bp after start (+) |
| QPLH2711.AG | CTCACCATCGAAAAGTCCAG | sgRNA target | 50 bp after start (+) |
| PBXV3009.AD | GTTTAAATTACGATATAAGA | sgRNA target | 19 bp before start (+) |
| PBXV3009.AE | GATGCCTCTGGACTTTTCGA | sgRNA target | 23 bp after start (+) |
| PBXV3009.AF | GGCCTTGGAGATCCTTCTCT | sgRNA target | 83 bp after start (-) |
| Hcyg_sens2_3F | GGAACCGTTTGCCACAGTAC | qPCR | Efficiency: 102% |
| Hcyg_sens2_3R | CTCTGCAGCAAGACATTCGG | qPCR | - |
| EF_AT_F1 | GCTGACGGTAAATGCCTCAT | qPCR | Efficiency: 99% |
| EF_AT_R1 | CAGGAGCGAACACAACAATG | qPCR | - |

**Table S6. CRISPR/Cas9 *sens-2* knockout efficiency estimates based on HRMs on egg DNA.**

| | | | | | Egg HRMs | | |
|---|---|---|---|---|---|---|---|
| **sgRNA 1** | **sgRNA 2** | **uM each sgRNA** | **uM Cas9** | **# Injected** | **Normal** | **Mutant** | **Predicted efficiency** |
| QPLH2711.AD | QPLH2711.AE | 0.2 | 0.2 | 30 | 13 | 6 | 32% |
| QPLH2711.AD | QPLH2711.AG | 0.2 | 0.2 | 31 | 9 | 13 | 59% |
| QPLH2711.AE | QPLH2711.AG | 0.2 | 0.2 | 31 | 9 | 4 | 31% |
| QPLH2711.AD | QPLH2711.AE | 1 | 1 | 19 | 15 | 1 | 7% |
| QPLH2711.AD | QPLH2711.AG | 1 | 1 | 28 | 3 | 3 | 50% |
| QPLH2711.AE | QPLH2711.AG | 1 | 1 | 28 | 9 | 4 | 44% |
| QPLH2711.AD | QPLH2711.AE | 5 | 5 | 25 | 2 | 7 | 78% |
| QPLH2711.AD | QPLH2711.AG | 5 | 5 | 24 | 6 | 6 | 50% |
| QPLH2711.AE | QPLH2711.AG | 5 | 5 | 24 | 0 | 5 | 100% |
| PBXV3009.AD | PBXV3009.AE | 0.2 | 0.2 | 67 | 14 | 2 | 13% |
| PBXV3009.AD | PBXV3009.AE | 1 | 1 | 55 | 8 | 3 | 27% |
| PBXV3009.AD | PBXV3009.AE | 5 | 5 | 48 | 6 | 2 | 25% |
| PBXV3009.AF | - | 0.2 | 0.2 | 37 | 12 | 4 | 25% |
| PBXV3009.AF | - | 1 | 1 | 21 | 10 | 1 | 9% |
| PBXV3009.AF | - | 5 | 5 | 33 | 7 | 1 | 13% |

**Table S7. CRISPR/Cas9 injections, survival, and mosaicism. Mixes were injected into *Heliconius cydno galanthus* or *H. c. alithea* (any color) - no differences in survival or mutation were observed between groups.**

| sgRNA1 | sgRNA2 | Concentration (uM) | Eggs injected | Larvae | Pupae | Mosaic |
|---|---|---|---|---|---|---|
| QPLH2711.AD | QPLH2711.AG | 0.2 | 374 | 14 | 5 | 1 |
| QPLH2711.AD | QPLH2711.AG | 1 | 1211 | 24 | 4 | 1 |
| QPLH2711.AD | QPLH2711.AE | 5 | 723 | 12 | 2 | 0 |
| QPLH2711.AE | QPLH2711.AG | 5 | 1372 | 18 | 6 | 2 |
